# Oncogenic transformation proceeds through a transient state of cellular plasticity constrained by lineage-specific barriers

**DOI:** 10.64898/2026.08.05.743116

**Authors:** Costakis Frangou, Alfiya Safina, Mairead Commane, Vidula Jahav, Ayah Salameh, Brian Buckley, Prashant Singh, Jesse Luke, Daniel Diaz, Katerina Leonova, Jianmin Wang, Katerina Gurova

## Abstract

Normal tissues frequently harbor oncogenic mutations without progressing to cancer, but the cellular basis of this resistance remains poorly defined. We asked whether transformation requires selection of a rare, permissive state within normal-like cell populations. Dermal fibroblasts and mammary epithelial cells responded uniformly to combined HRAS-G12V expression and p53 disruption. Cellular barcoding revealed no loss or enrichment of clones during morphological transformation, arguing against clonal selection as the primary driver of neoplastic reprogramming. Instead, all transduced cells underwent an early, shared transcriptional transition characterized by loss of differentiation markers, induction of RAS-associated and inflammatory programs, activation of alternative-lineage signatures, increased single-cell entropy, and chromatin decondensation. These changes were transient: entropy and chromatin accessibility subsequently declined, and some cells moved toward the control transcriptional state, whereas others stabilized in altered states. The two lineages followed distinct trajectories. Fibroblasts showed greater initial transcriptional plasticity but subsequently reverted more strongly toward the normal state, whereas epithelial cells changed more gradually and continued to diverge from it, suggesting a stronger barrier to transformation in the mesenchymal lineage. Thus, oncogenic perturbation initiated reprogramming throughout the population but did not uniformly produce a stable transformed state. Together, these findings support a model in which normal-like cells tolerate oncogenic mutations not because most cells fail to respond, but because a p53-independent, cell-intrinsic barrier limits the stabilization of malignant transformation following a transient period of heightened plasticity. This framework may facilitate the identification of mechanisms that constrain tumor initiation..

**Significance Statement:** Loss of TP53 and activation of oncogenic RAS are common drivers of human cancer and frequently coexist in the same tumor. Nevertheless, these alterations often fail to induce malignant transformation. To investigate why, we introduced both alterations into fibroblasts and epithelial cells and tracked their responses using clonal barcoding, single-cell transcriptomics, and phenotypic assays. Initially, cells uniformly reprogrammed their transcriptional and chromatin states without detectable clonal selection. They then diverged: some reacquired transcriptional profiles resembling those of the original normal cells, whereas others became transformed. Thus, normal cells resist transformation through an intrinsic barrier that operates despite p53 disruption. Because this transition is transient and reversible, it provides a tractable window for studying the earliest stages of cancer development.

## Introduction

Cancer arises through the transformation of normal cells into malignant cells, a process driven by genetic, epigenetic, and microenvironmental factors (1, 2). The gene-centric model of tumorigenesis posits that mutations in tumor suppressor genes or proto-oncogenes directly induce transformation. This model is supported by the recurrence of these mutations across patient tumors and by experimental validation in genetically engineered mouse models (GEMMs). However, accumulating evidence indicates that genetic mutations alone are often insufficient for malignant transformation. For example, driver mutations have been detected in histologically normal tissues from aging individuals (3, 4), and GEMMs rarely develop the full potential tumor burden even when every cell in a tissue carries the same engineered driver mutations. The acquisition of additional somatic alterations typically accelerates tumor onset but does not substantially increase tumor number in GEMMs (5), (6). These observations point to intrinsic cellular barriers that limit malignant transformation.

Tumor suppression can arise from cell-intrinsic mechanisms, including chromatin regulation, transcriptional reprogramming, and metabolic checkpoints, as well as from environmentally driven mechanisms involving signals from stromal, immune, or tissue-niche compartments. Environmental factors are powerful modulators of tumor initiation and progression, and their roles and underlying mechanisms have been demonstrated in multiple studies (reviewed in ref. 9). Several mechanisms of cell-intrinsic resistance to malignant transformation have also been identified, most notably p53 activation in response to oncogene activation (7, 8). However, nonmutational, cell-intrinsic barriers to transformation remain poorly understood.

Epigenetic transitions associated with processes such as inflammation and wound healing can alter cellular sensitivity to oncogenic transformation (1–5). We therefore propose that, under basal conditions, rare transitional epigenetic states may exist that render cells permissive to oncogenic transformation. Alternatively, the observation that mutated cells in normal tissues frequently occur as clones (3, 4) suggests that many cells may initiate the transformation process but are subsequently restrained by as-yet-unknown barriers that restore cellular homeostasis despite the acquisition of oncogenic mutations (8, 10–12). We therefore asked whether a specific preexisting epigenetic state sensitizes cells to oncogene-induced transcriptional reprogramming or whether additional intrinsic barriers can return cells to a normal state despite the acquisition of oncogenic mutations.

To address this question, we developed an in vitro experimental system to compare cell-autonomous responses to defined oncogenic perturbations across distinct human cell types. We combined single-cell transcriptomics to capture dynamic changes in gene expression, cellular barcoding to track clonal behavior, and phenotypic assays to measure tumorigenic potential. We applied this approach to immortalized, nontransformed human fibroblasts and epithelial cells perturbed with HRASG12V and dominant-negative p53, a canonical oncogenic pair that frequently co-occurs in tumors. A uniform early response shared across cell types resolved into lineage-specific trajectories: a substantial proportion of cells returned to a near-normal state, whereas others continued toward a fully transformed phenotype. These findings suggest the existence of p53-independent, cell-intrinsic mechanisms that resist malignant transformation.

## Results

### Establishing Experimental Conditions for Modeling Transformation in Human Cells

To investigate oncogene-driven reprogramming across tissue contexts, we used three normal-like, genetically stable human cell models: primary normal dermal fibroblasts (NDFs), immortalized mammary epithelial cells (MCF10A) (13), and kidney epithelial cells (NKEs) (14).

We used lentiviral transduction to introduce a bicistronic cassette encoding the dominant-negative p53 fragment GSE56, which contains part of the p53 tetramerization domain, and HRASG12V, linked by an internal ribosome entry site (IRES). We refer to this construct as GR. HRASG12V constitutively activates RAS signaling (15), whereas GSE56 inhibits p53 tetramerization, leading to the stabilization of inactive monomeric p53 (16, 17). To preserve the full spectrum of cellular and molecular responses we excluded artificial antibiotic selection against nontransduced cells, since this selection can also eliminate cells unable to transform. We adjusted the multiplicity of infection (MOI) for each cell type to achieve nearly ubiquitous transgene delivery. Sustained cassette expression was confirmed by p53 stabilization in GR-transduced cells, whereas wild-type p53 was undetectable in controls (Fig. S1 A–D). Cells were sampled at multiple time points after transduction in several independent experiments. Findings reproduced across experiments are presented in the main figures, and examples from replicate experiments are presented in the supplementary data.

Oncogene expression induced cell type–specific changes in growth and morphology. GR-transduced epithelial cells grew more slowly than empty-vector (EV) controls, whereas GR-transduced fibroblasts proliferated more rapidly (Fig. 1 A–C). By day 4, MCF10A cells had transitioned from compact epithelial sheets to elongated, spindle-shaped cells consistent with epithelial–mesenchymal transition (EMT), whereas NDF-GR cells became larger and flatter and developed epithelial-like regions suggestive of mesenchymal–epithelial transition. Both cell types contained numerous stressed cells characterized by reduced spreading, increased refractility, partial rounding, and cytoplasmic shrinkage relative to untransduced or EV-transduced cells. These stressed cells disappeared and cell morphology stabilized within 2 to 3 weeks (Fig. 1 D and E).

**Figure 1.**
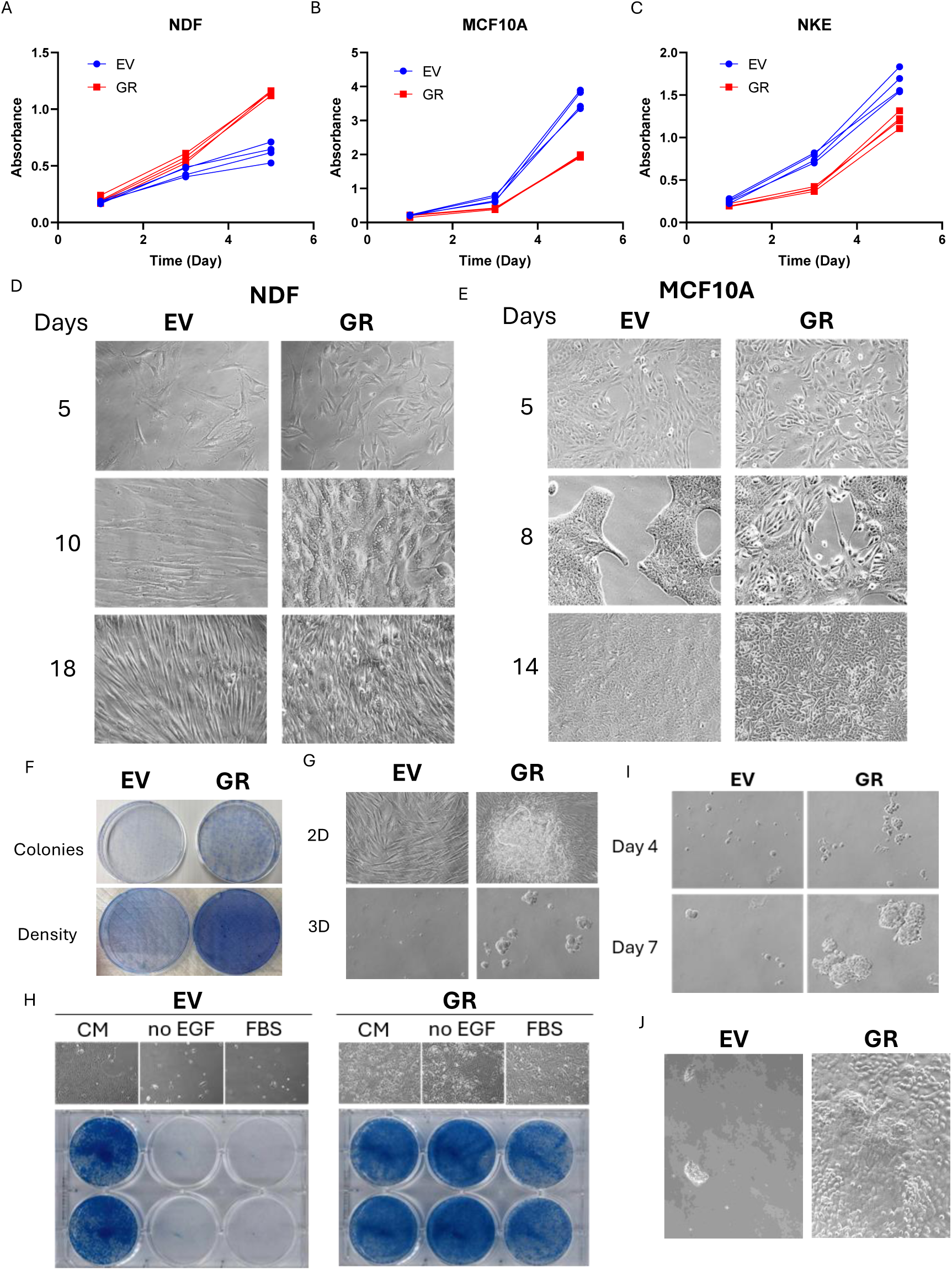
Phenotypic changes of cells upon GR transduction. A-C. Growth of NDF (A), MCF10A (B) and NKE (C) cells plated at 72 hours after transduction with GR or EV viruses. At each time point, replicate plates of cells were fixed and stained with methylene blue. Y-axis – absorbance of methylene blue extracted from stained plates. Three biological replicates are shown. D, E. Photographs of NDF (D)and MCF10A (E) cells at different time points after EV or GR transduction. F. Growth of EV or GR transduced MCF10A cells in media of different compositions. CM - complete medium. Photographs of cells and methylene blue stained plates (two replicate wells). G. Growth of EV and GR transduced MCF10A cells in the semisolid medium at days 4 and 7 after plating. H. Methylene blue stained plates of EV or GR transduced NDF cells plated at low density for colony formation (upper row) or at higher density for the observation of the effect of contact inhibition (bottom row). I. Colony formation of EV and GR transduced NDF cells in 2D and 3D conditions. Upper row – growth as a monolayer in 2D with a focus of cells growing on top of monolayer in case GR. Bottom row – growth of cells in semisolid medium. J. Photographs of EV or GR transduced MCF10A cells plated into standard 2D conditions after growing in 3D.

We next investigated whether GR cells had acquired functional hallmarks of transformation: anchorage-independent growth, growth-factor independence, and loss of contact inhibition. NDF-GR fibroblasts formed colonies from single cells and showed reduced contact inhibition (Fig. 1F). In semisolid medium, NDF-GR cells formed large, irregular colonies, whereas NDF controls formed none (Fig. 1G). MCF10A-GR cells proliferated without EGF and tolerated fetal bovine serum (FBS), conditions that suppress the growth of parental cells (Fig. 1H). In semisolid medium, MCF10A-GR cells formed large, irregular colonies, whereas parental MCF10A cells formed small, round colonies (Fig. 1I). When transferred from semisolid medium to two-dimensional culture, GR-MCF10A cells reattached within 24 h and continued to proliferate. Only a few EV cells reattached, and these cells failed to expand (Fig. 1J). In contrast, NKE cells showed no morphological differences following GR transduction and were excluded from downstream molecular analyses (data not shown).

### Clonal Architecture of the Early Transformation Response

To determine whether the transformed phenotype arose through selection of a rare, permissive subpopulation or through a uniform transition of the entire population, we used cellular barcoding to track clonal dynamics over time. We observed no reduction in the number of barcodes during transformation (∼15 to 20 days; Fig. 2 A, B and Fig. S2A). We also measured the Shannon entropy of the barcode distribution, which decreases when a population becomes dominated by a few clones (18). During transformation, entropy remained constant in both NDF and MCF10A populations (Fig. 2 C, D), indicating no detectable clonal selection. Rarefaction curves plateaued below the sequencing depths used, confirming saturated sampling in both lineages (Fig. 2 E, F). Clonal complexity declined only after prolonged passaging; however, this decline did not differ between EV and GR populations, indicating random clonal drift during extended passage. Similar barcodes were also enriched across replicate experiments at later time points, but this enrichment did not differ between EV and GR cells, suggesting that GR conferred no fitness advantage beyond random differences among cells within each population (Fig. S2 C, D). Thus, the early transformation response was population-wide rather than driven by rare preexisting clones.

**Figure 2.**
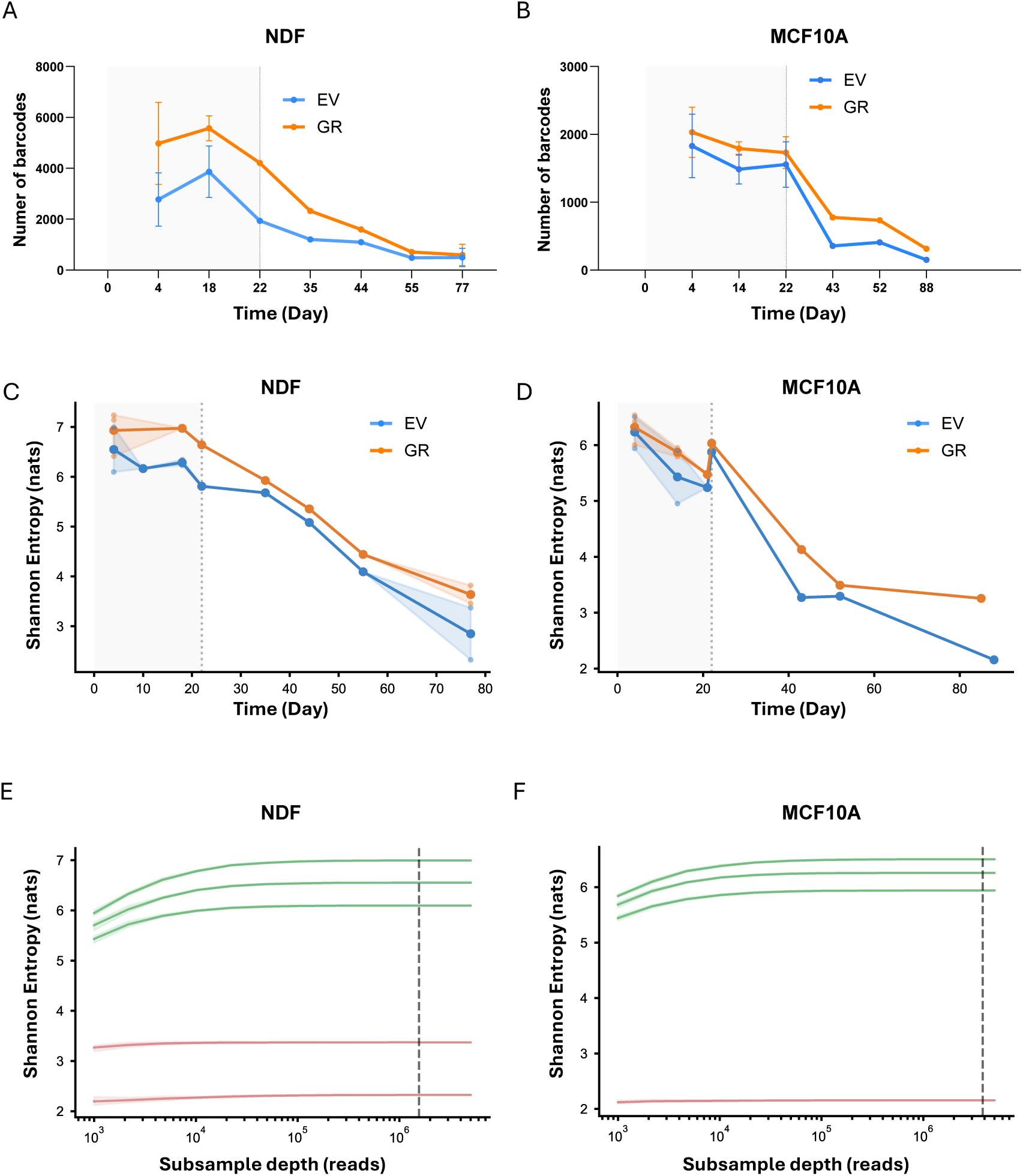
Population-scale clonal complexity of barcoded NDF and MCF10A populations during transformation. A, B. Quantitation of changes in the number of detected barcodes in NDF (A) or MCF10A (B) cell populations at different time points after transduction of EV or GR viruses. Mean of two or three independent experiments +/- SD. The grey background indicates window of morphological transformation from day 0 to day 22, with a dotted line at the day-22 boundary. C, D. Shannon entropy of barcode distributions in NDF (C; n = 23 samples) and MCF10A (D; n = 20 samples) populations at different time points after transduction of EV or GR viruses. Mean values and the minimum and maximum across biological replicates (shaded ribbon). The grey background is the same as in panels A and B. E, F. Rarefaction analysis for NDF (E) and MCF10A (F). Shannon entropy of the barcode distribution against subsampled read depth for one sample. Color marks the two sampling windows: 0 – 22 days - samples in green, after 22 days - samples in red. The dashed vertical line marks the rarefaction depth applied downstream. All curves plateau below that depth, confirming saturated sampling in both lineages.

### Nonlinear Transcriptome Changes during Malignant Transformation

Because the transformation response was not limited to the selection of preexisting clones, we profiled gene-expression changes over time by measuring mRNA abundance at single-cell resolution using SPLiT-seq (Parse Biosciences). We analyzed NDF and MCF10A cells that were untransduced, EV-transduced, or GR-transduced and sampled at multiple time points during transformation. SPLiT-seq indexes fixed cells through iterative barcoding, enabling the multiplexing of time points and conditions within a single library. This strategy yielded single-cell datasets for each experiment (Fig. 3 A, B and Table S1).

**Figure 3.**
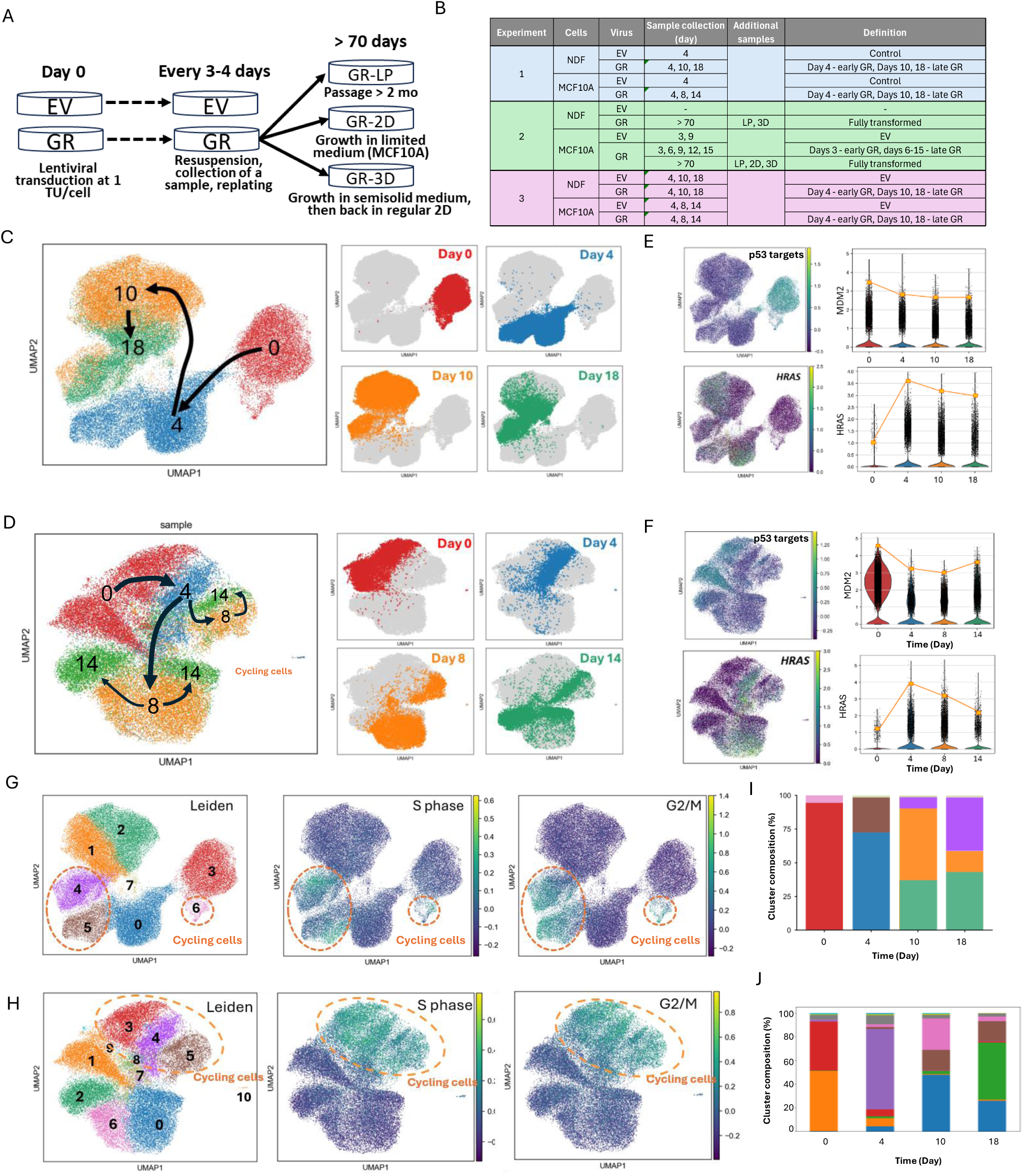
Non-linear transcriptome changes during oncogenic transformation of NDF and MCF10A cells. A. Scheme of experiment and sample collection for scRNA-seq. B. List of all samples used for scRNA-seq in different experiments. C, D. UMAP visualization of single-cell transcriptomes of NDF (C) and MCF10A (D), with individual cells colored according to sample name. Numbers indicate day after transduction. 0 – cells transduction with EV. Large plots on the left show all samples together. Small plots on the right show cells of individual samples, with all other cells in gray. E, F. Dynamic of expression of p53 target genes and HRAS. UMAP feature plots showing log-normalized expression of the indicated genes. Each point represents one cell; color indicates expression level. Violin plot shows distribution of expression of a gene between all cells in a sample. Orange lines connect 90th percentile of corresponding gene(s) expression. G, H. UMAP plots with Leiden (left) clusters of NDF (G) and MCF10A (H) cells and the presence of gene markers of S (central) or G2/M (right) phases of cell cycle. Cells with cell cycle markers are shown as orange dotted circles on each plot. I, J. Distribution of Leiden clusters between samples of NDF (I) or MCF10A (J) cells collected at different times.

Fibroblasts and epithelial cells occupied distinct transcriptional spaces in every replicate (Fig. S3A); we therefore analyzed each cell line separately. p53 inactivation and HRASG12V expression were confirmed at the transcript level: GR cells showed reduced expression of canonical p53 target genes and increased total HRAS mRNA, which was almost undetectable in control cells. (Fig. 3 C–F; Figs. S3 B–E, S4 A–C, and S5 A–C). Notably, HRAS expression peaked early and then declined, although it remained above the levels observed in EV controls. This pattern suggests selective constraints on high RAS expression even when p53 is nonfunctional

Unbiased, sample-agnostic clustering visualized by UMAP revealed three well-separated groups of fibroblasts: control, early GR (day 4), and late GR (days 10 and 18), whereas epithelial cells did not form clearly separated groups (Fig. 3 C, D and Figs. S4A, S5A). This finding suggests that transcriptional changes during transformation are stronger or more uniform in mesenchymal cells than in epithelial cells. Based on cell-cycle markers, the transcriptomes of both cycling and noncycling cells differed among control, early-GR, and late-GR populations (Fig. 3 G, H and Figs. S4D, S5D), indicating that the effects of GR transduction were independent of cell-cycle state.

In both NDF and MCF10A cells, the earliest GR samples (days 3 to 4) segregated from EV controls on UMAP plots, indicating a rapid and relatively uniform initial transcriptional response to GR. At later time points (days 10 to 18 for NDF and days 6 to 15 for MCF10A), GR cells shifted further from the early-GR state but substantially overlapped across adjacent time points, suggesting continuous transcriptional progression rather than discrete state transitions. Within these late populations, a subset of GR cells in each lineage moved toward, but did not fully converge with, control profiles, whereas other cells diverged further (Fig. 3 C and D and Figs. S4A and S5A).

The late-stage populations could represent either discrete states or a continuum. To distinguish between these possibilities, we analyzed Leiden clusters within each population. NDF cells resolved into eight clusters that diversified at late time points (Fig. 3 G and I and Fig. S4D), whereas MCF10A cells resolved into 10 clusters whose identities shifted progressively over time (Fig. 3 H and J and Fig. S5D). Cluster proportions over time also differed substantially between the lineages (Fig. 3 I and J). To assess cluster discreteness, we applied partition-based graph abstraction (PAGA; (19)), which weights connections between clusters according to transcriptomic similarity, such that denser edges indicate continuous transitions and sparser edges indicate discrete boundaries. NDF clusters were more sparsely connected than MCF10A clusters (Fig. S3 F and G), indicating that fibroblasts pass through discrete states, whereas epithelial cells change more continuously. Moreover, trajectories in both lineages were nonlinear and divergent, with similar proportions of cells in late-GR populations moving closer to the control state (cluster 2; Fig. 3 G, H). These observations were confirmed in an independent experiment (NDF cluster 1 and MCF10A cluster 0 on Figs. S4D and S5D respectively).

### Loss and Recovery of Differentiation Markers during Transformation

To characterize changes in transcriptional programs during transformation, we performed gene set enrichment analysis (GSEA) on genes differentially expressed between time points. We grouped gene sets with similar annotations into broader categories and organized the results of this analysis as circular barplots, explained on Fig. S8A. The predominant category in control fibroblasts comprised markers of the matrisome, collagen organization and remodeling, and the extracellular matrix (ECM); we designated this category “Fibroblast Identity” (Fig. 4 A, B). Control NDFs were also enriched for genes that are downregulated in cancer and inflammation. Control epithelial cells were enriched for p53 target genes and markers of epithelial differentiation, including sterol-metabolism genes and keratins (Fig. 5 A, B). Both cell types followed a common trend: early-GR cells differed most strongly from control cells: they lost a substantial proportion of their respective fibroblast or epithelial identities, whereas later GR cells partially recovered these identities (Figs. 4A and 5A). This trend illustrated with several individual fibroblast-or epithelial-specific genes (Figs. 4C and 5C). Over-representation analysis (ORA; Enrichr, MSigDB C2; (20)) quantitatively confirmed these observations (Figs. S6 and S7).

**Figure 4.**
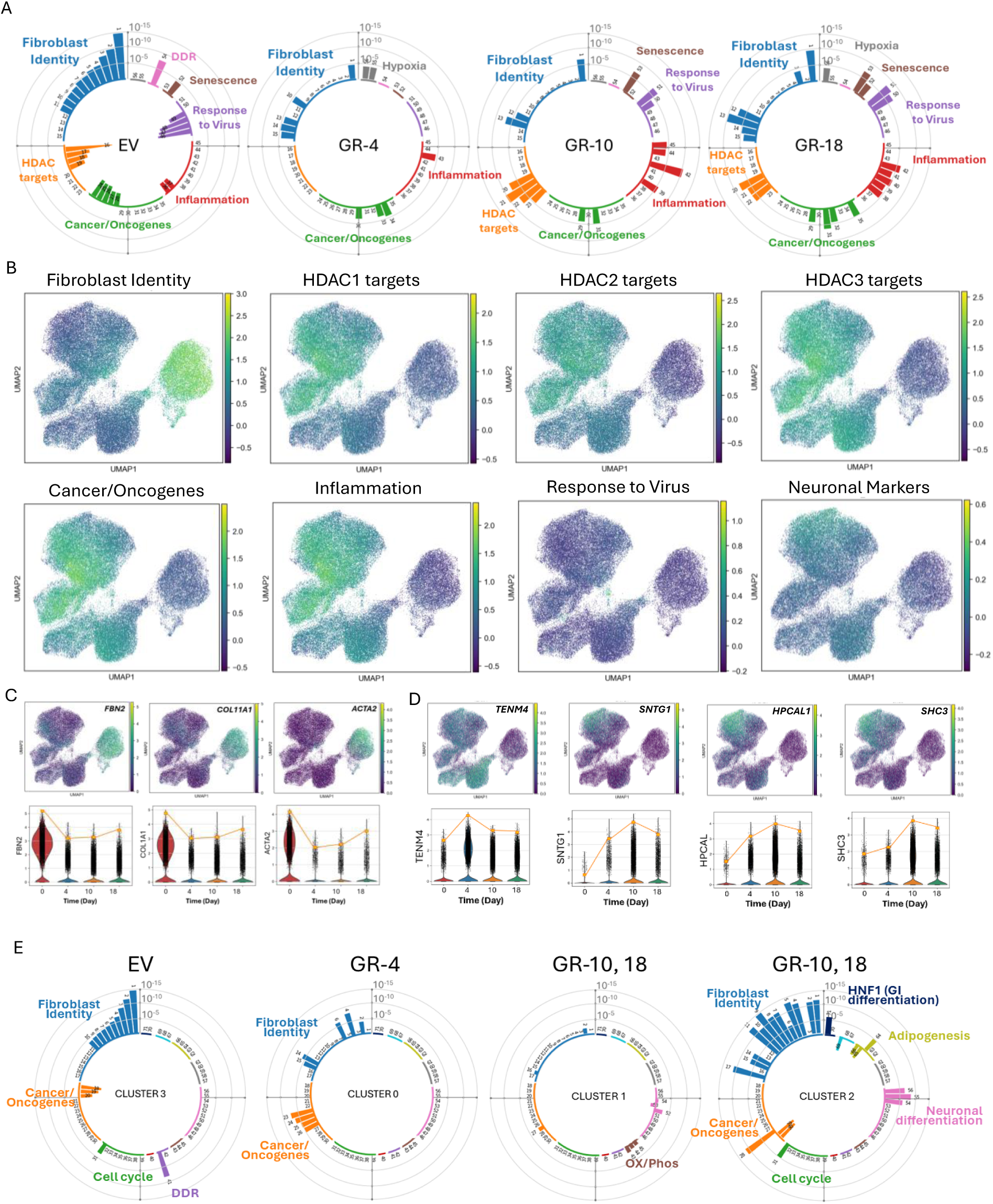
Transcriptional changes during transformation of NDF cells. A. Circular barplots demonstrating presence of transcriptional signatures in population of NDF cells at different time points after transduction of EV or GR viruses. Bars are q-values of enrichment of individual gene sets organized by categories in different groups and colors. Color corresponds to the colored category name. Number are gene lists shown in table S2. Bars looking outside of a circle are upregulated genes, bars looking inside the circle are downregulated genes. B. Presence of several transcriptional signatures in NDF cells shown on UMAP plots and color-coded according to the score of a signature. The same UMAP with sample names are shown on Fig. 3C. C. Examples of “Fibroblast Identity” genes. UMAP and violin plots showing log-normalized expression of the indicated genes. D. Neuronal markers expression in NDF cells. UMAP and violin plots showing log-normalized expression of the indicated genes. E. Circular barplots demonstrating presence of transcriptional signatures in population of NDF cells in different Leiden clusters and samples. Only clusters of non-cycling cells are shown. Cycling cluster are presented on Fig. S8A. Circular barplots are designed as described in A.

**Figure 5.**
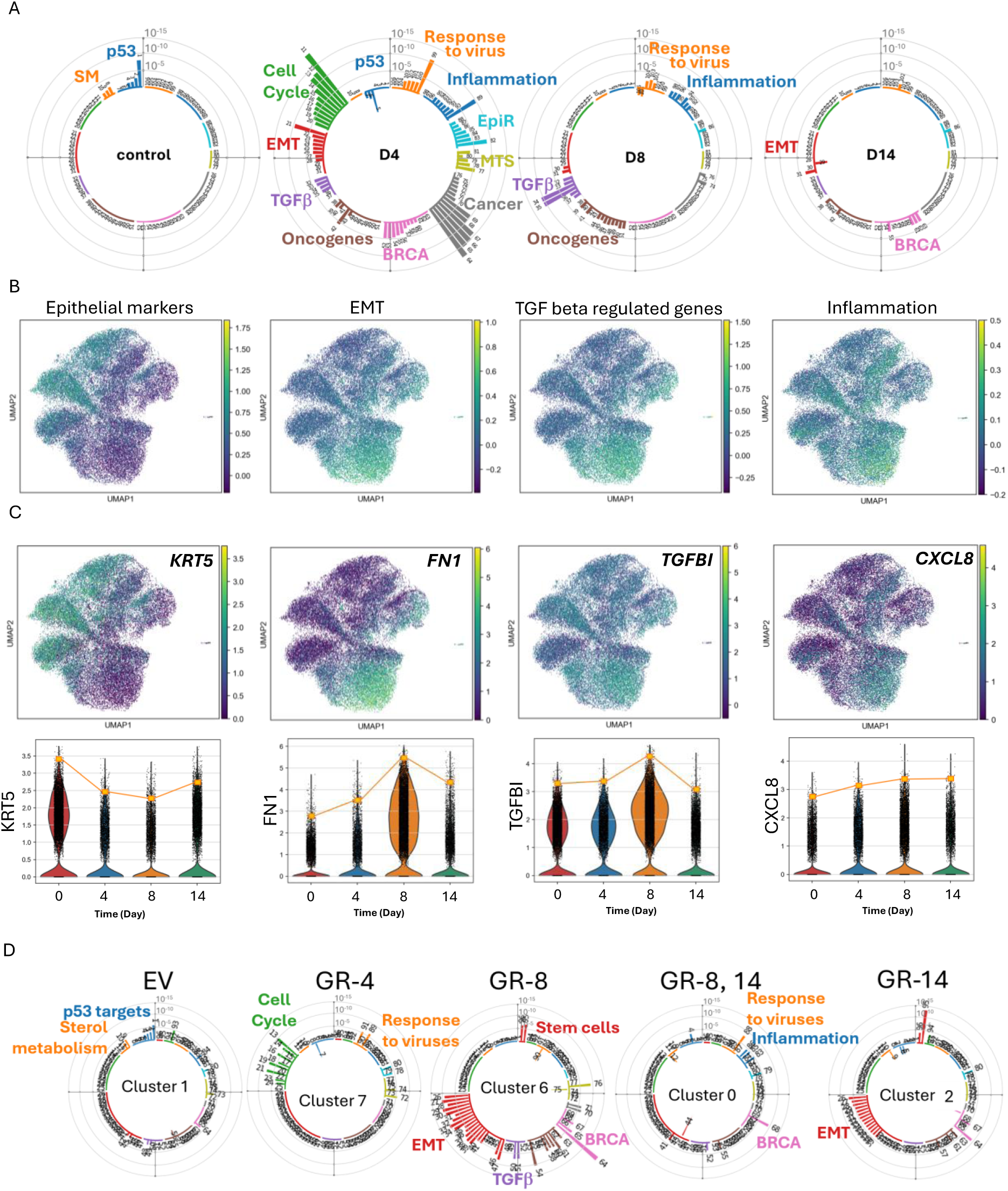
Transcriptional changes during transformation of MCF10A cells. A. Circular barplots demonstrating presence of transcriptional signatures in population of MCF1-A cells at different time points after transduction of EV or GR viruses. Bars are q-values of enrichment of individual gene sets organized by categories in different groups and colors. Color corresponds to the colored category name. Number are gene lists shown in table S3. Bars looking outside of a circle are upregulated genes, bars looking inside the circle are downregulated genes. B. Presence of several transcriptional signatures in MCF10A cells shown on UMAP plots and color-coded according to the score of a signature. The same UMAP with sample names are shown on Fig. 3D. C. Examples of genes, representing “Epithelial Markers” (KRT5), EMT (FN1), “TGF Regulated Genes” (TGFBI) or “Inflammation” (CXCL8). UMAP and violin plots showing log-normalized expression of the indicated genes. D Circular barplots demonstrating presence of transcriptional signatures in population of MCF10A cells in different Leiden clusters and samples. Only clusters of non-cycling cells are shown. Cycling cluster are presented on Fig. S8B. Circular barplots are designed as described in A.

Early-GR fibroblasts (day 4) had the fewest significantly enriched pathways (Fig. 4A). By day 10, GR cells had upregulated genes associated with “Inflammation” and “Response to Viruses.” These cells also upregulated genes that are inhibited by various histone deacetylases (HDACs) (Fig. 4 A–C and Fig. S6). NDF-GR cells initially increased and later subsequently reduced the expression of alternative-lineage markers, including neuronal markers (Fig. 4 B and D and Fig. S6).

Unlike fibroblasts, early-GR epithelial cells had the largest number of enriched pathways. These included cell-cycle programs; EMT-related programs, TGFβ-regulated genes, integrin signaling; oncogenic signatures, including several gene sets enriched in breast and other cancers; and metastasis-related programs. On days 3 to 4, GR epithelial cells also showed significantly increased expression of genes associated with epigenetic regulation and inflammation. At later times, only a few of these signatures persisted, including transformation, EMT, and inflammation, and their enrichment was less significant (Fig. 5 A and B and Fig. S7).

Leiden clustering showed that control fibroblasts and epithelial cells each comprised two clusters: cycling and noncycling cells. Early-GR NDFs (day 4) retained the same two clusters, whereas most early-GR MCF10A cells formed one large cluster of cycling cells and several smaller clusters shared with other samples (Fig. 3 G–J, Fig. 4E, and Fig. 5D). The two major clusters of noncycling late-GR fibroblasts differed markedly from each other (clusters 1 and 2; Fig. 4E). Cluster 1 had almost no enriched pathways, except for modest enrichment of oxidative phosphorylation genes based on P values. Cluster 2 fibroblasts regained high “Fibroblast Identity” scores that approached those of control cells and showed reduced representation of “Cancer-Related” gene sets; some of these sets were downregulated, as in control cells. Unlike control cells, however, cluster 2 cells expressed markers of other lineages, including neuronal, mammary-gland, and adipocyte markers (Fig. 4E).

The principal difference among Leiden clusters of noncycling GR epithelial cells was the presence or absence of an EMT signature. Control cells completely lacked this signature, whereas two of the three major GR clusters (3 and 6) contained numerous EMT- and TGFβ-regulated gene sets. GR cluster 0 lacked an EMT signature and had minimal representation of other signatures, resembling cluster 1 of NDF cells (Fig. 5D). Markers of alternative differentiation, including adipocyte and neuronal differentiation, also emerged in epithelial cells; neuronal markers were especially prominent in clusters of cycling GR cells (Fig. S8B, C).

We observed enrichment of two classes of proinflammatory genes: one comprising NF-κB-related pathways and the other comprising the response to viruses, including interferon (IFN)-responsive genes. Because these classes overlap substantially, we examined individual genes known to be regulated predominantly by NF-κB or IFN. In NDF cells, NF-κB-regulated genes were elevated almost exclusively on days 10 and 18 and showed considerable heterogeneity (compare CXCL1 and IL1B; Fig. S9A). In MCF10A cells, these genes were elevated almost exclusively on day 8 (Fig. S10A). In contrast, IFN-responsive genes were upregulated relatively uniformly in NDF Leiden cluster 7, which contained GR cells from multiple time points. In MCF10A cells, a small proportion of cells with expression of IFN-responsive genes was dispersed across multiple groups (Figs. S9B and S10B).

These observations can be summarized as follows. (i) The earliest and most pronounced change during oncogene-induced transformation was reduced or lost expression of lineage-specific genes. This change was accompanied by (ii) overexpression of multiple genes associated with the RAS signature and other cancer-related processes, (iii) overexpression of genes normally silenced by HDACs, and (iv) heterogeneous expression of inflammation-related genes. (v) Later in transformation, some cells continued to lose their identity and showed no significantly enriched pathways among group-specific genes. In contrast, another group restored the expression of some differentiation-specific genes but also began to express markers of alternative lineages, including neuronal, adipose, and mammary-gland markers in fibroblasts and EMT markers in epithelial cells.

### Chromatin Decondensation during Transformation

The enrichment of HDAC-target genes and other transcriptional changes—including loss of lineage identity, emergence of alternative-lineage signatures, and activation of inflammatory and interferon responses—suggested that cells passed through a less lineage-committed, more plastic state. To quantify this plasticity at single-cell resolution, we calculated two entropy measures across the time course: Correlation of Connectome and Transcriptome (CCAT), a network-based proxy for signaling entropy (21), and per-cell Shannon entropy. Both measures increase as cells lose lineage commitment. Relative to matched EV controls, GR cells showed transient increases in both measures, which peaked at day 4 in NDF and MCF10A cells and subsequently resolved. At the peak, the GR-minus-EV difference reached +0.015 in NDF and +0.018 in MCF10A cells for network entropy (Fig. 6A) and +0.20 and +0.33, respectively, for Shannon entropy (Fig. 6B). At later time points, these measures returned to EV levels in NDF cells and fell below EV levels in the other conditions. Thus, transformation began with a brief period of increased transcriptional plasticity that was most pronounced in the epithelial lineage before cells settled into stable states.

**Figure 6.**
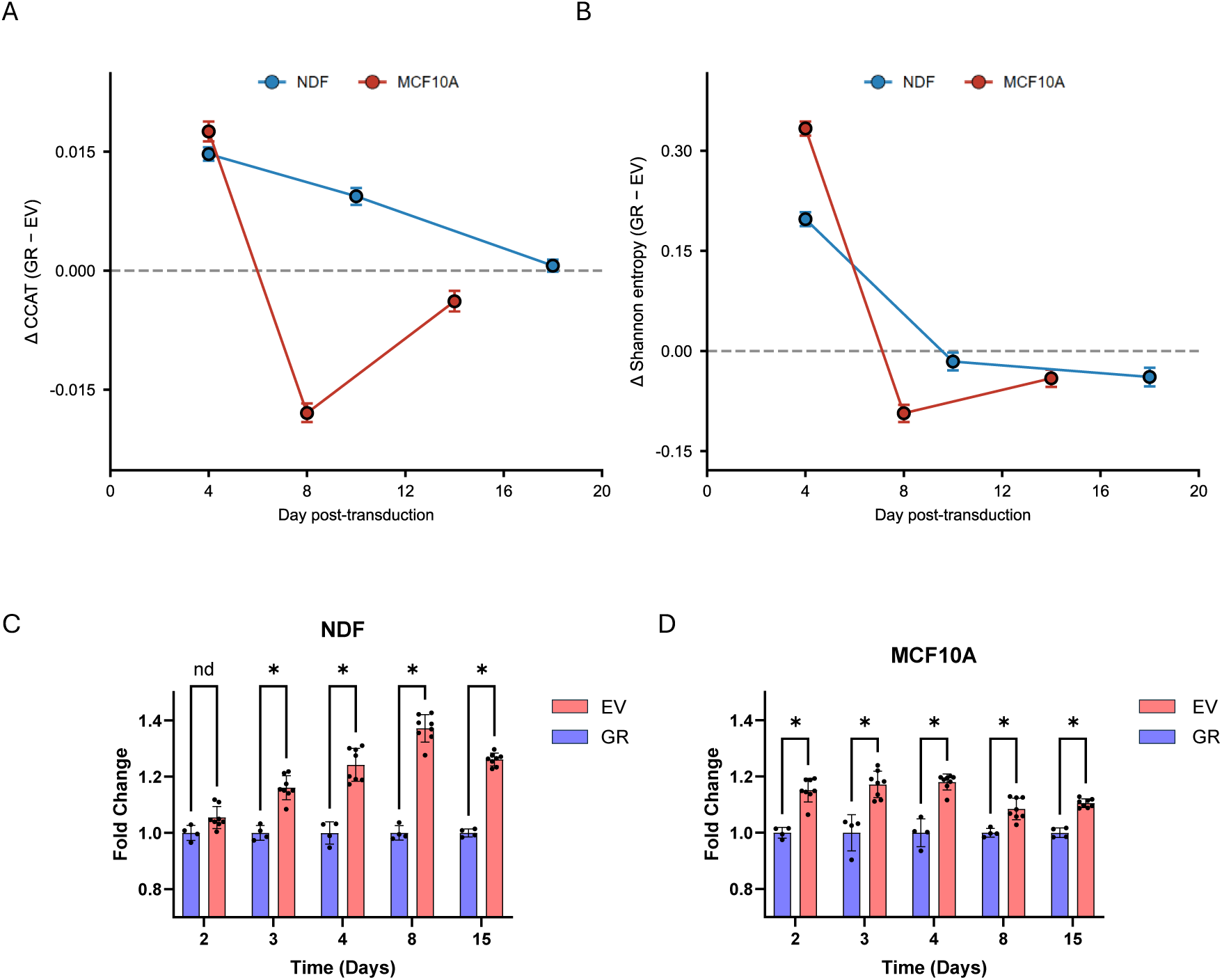
Transient increase in single-cell entropy and subsequent genome-wide chromatin decondensation after GR transduction in fibroblast and epithelial cells. A, B. Per-cell entropy relative to the EV control over the transformation time course, in the one experiment with a matched GR and EV time course per lineage (the third transduction experiment). A. Network entropy (CCAT, a network-based proxy for signaling entropy). B. Per-cell Shannon entropy. Each panel plots the GR-minus-EV difference at each day for NDF (blue) and MCF10A (red); points show the difference in means, error bars the 95 percent bootstrap interval over cells, and the dashed line marks zero; points connected in day order. The cell is the unit of analysis, so these are descriptive effect sizes rather than replicate-level inference. Cells per day (GR, EV): NDF 7,133 and 3,696 at day 4, 3,043 and 3,081 at day 10, 3,730 and 4,185 at day 18; MCF10A 3,915 and 3,789 at day 4, 4,304 and 3,753 at day 8, 4,097 and 3,094 at day 14. C, D. Chromatin accessibility in NDF (C) and MCF10A (D) cells over the time course, measured by fluorescent DNA-binding ligand staining of fixed cells (nuclear fluorescence intensity, arbitrary units). Bars are mean accessibility of four (EV) or eight (GR) biological replicates including two sets of independent virus preparation. Asterisks are p-values < 0.05 of multiple paired T-tests.

To validate these findings experimentally, we measured chromatin accessibility directly using a fluorescent DNA-binding ligand that preferentially labels nucleosome-free DNA (22). We quantified nuclear fluorescence intensity in GR- and EV-transduced NDF and MCF10A cells at multiple time points after transduction. As predicted, chromatin accessibility increased in both lineages following GR transduction, but with distinct kinetics. In NDF cells, accessibility showed a delayed peak at day 8, consistent with the later phenotypic response of fibroblasts (Fig. 6C). In MCF10A cells, accessibility peaked earlier, on days 3 to 4, coinciding with rapid morphological changes in these cells. Accessibility subsequently declined partially but remained above EV levels (Fig. 6D). Thus, transformation was characterized by transient chromatin decondensation whose timing paralleled the phenotypic response of each cell line, suggesting lineage-specific kinetics. Together, the single-cell entropy and direct chromatin-accessibility measurements identify a transient, self-limiting period of heightened plasticity at the onset of transformation. This period begins as cells depart from their initial state and subsides as lineage-specific programs take hold.

### Transcriptional States after Selection for Transformed Phenotypes

Both fibroblasts and epithelial cells reached a fully transformed state within 2.5 weeks, as indicated by anchorage-independent growth, growth-factor independence, loss of contact inhibition, and stable morphological state. However, transcriptional trajectories showed that a subset of cells drifted back toward control profiles by the end of this period, leaving the stability of the transformed state unresolved. We therefore examined the properties acquired by GR cells after additional selection for a transformed phenotype, either through maintenance in the absence of growth factors (GR-2D) or growth in semisolid medium (GR-3D; see Materials and Methods for details). In parallel, we collected cells maintained under standard conditions for more than 2 months (“long-passaged cells,” GR-LP). After selection, all cells were returned to identical standard growth conditions for at least 2 weeks before SPLiT-seq analysis (Fig. 3B). We collectively refer to these populations as cells selected for the transformed phenotype (STP).

To identify transcriptional differences between selected and preselection states, we compared STP populations with GR cells at late transformation time points (late GR; days 10 to 18 for NDF and days 8 to 15 for MCF10A). NDF STP fibroblasts formed an isolated UMAP cluster distinct from both late-GR and EV cells and positioned approximately equidistant from them (Fig. 7A). By contrast, STP MCF10A cells shifted away from EV cells, beyond the late-GR cells (Fig. 7C). Relative to late-GR cells, NDF STP cells partially restored the p53-target module (difference in mean score, +0.16; 95% CI, +0.15 to +0.16) and showed a modest reduction in HRAS expression (−0.12; 95% CI, −0.13 to −0.11; Fig. 7 B, E, and K). Conversely, STP MCF10A cells showed further reductions in p53-target expression and increases in HRAS expression (Fig. 7 D, F). STP fibroblasts also reexpressed the “Fibroblast Identity” module and showed reduced expression of alternative-lineage markers, including neuronal markers (Fig. 7 G and K), whereas epithelial cells showed further loss of epithelial markers and increased EMT/TGF-β signatures (Fig. 7 H and K). HDAC-target module scores decreased in fibroblasts but remained stable in epithelial cells (Fig. 7K). Consistent with these changes, the “Inflammation” signature decreased in fibroblasts but increased in epithelial cells. In contrast to this, the “Response to Viruses,” or IFN-response, signature became strongly activated in fibroblasts but was nearly undetectable in epithelial cells (Fig. 7 I–K).

**Figure 7.**
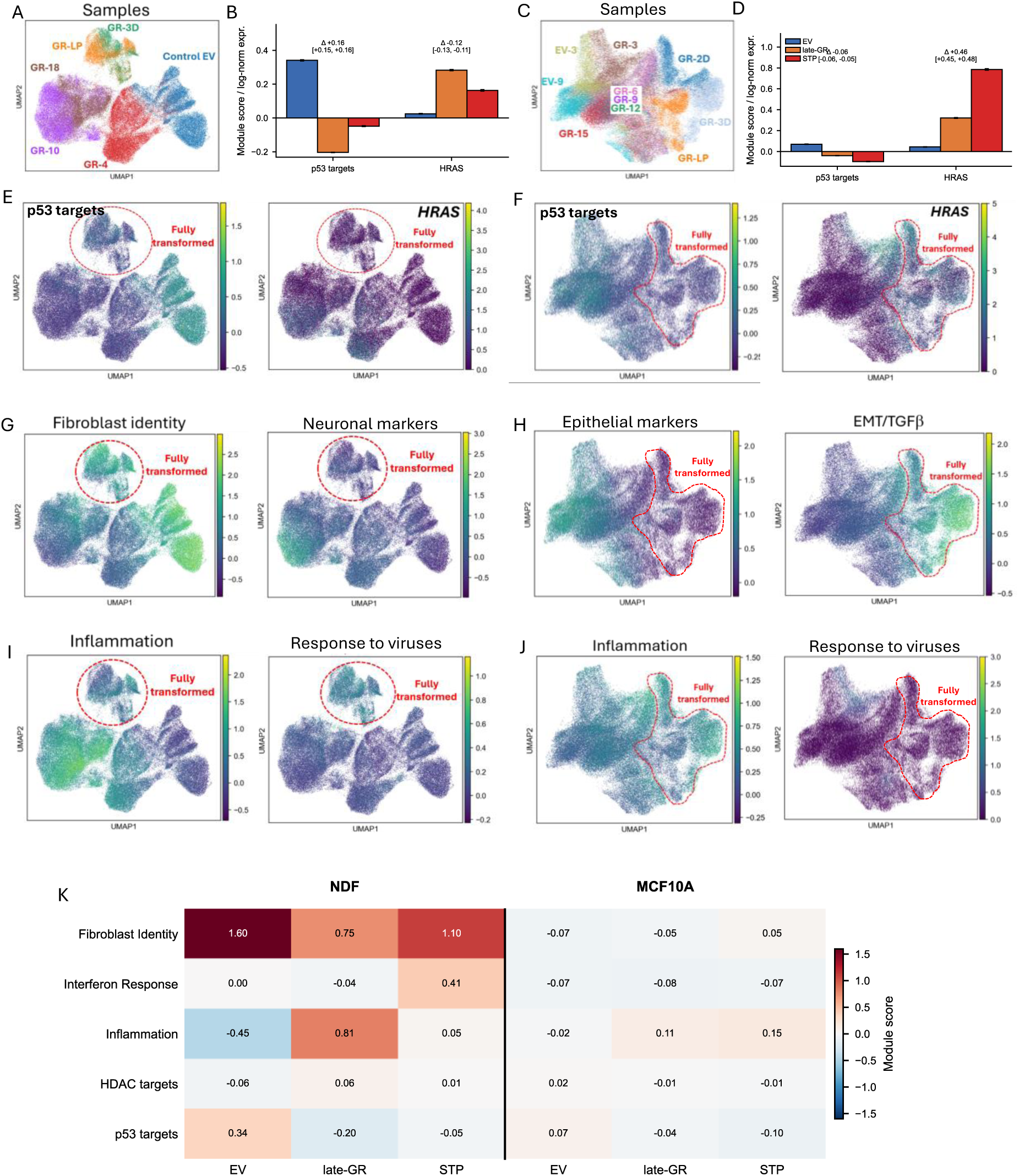
Distinct, lineage-specific stable states of selected transformed-phenotype (STP) populations in NDF and MCF10A cells. Conditions: EV (empty vector control), late-GR (GR at the late transformation time points, NDF days 10 to 18 and MCF10A days 6 to 15), and STP (selected transformed phenotype). A.C. UMAP feature plots visualizing distribution of individual NDF (A) and MCF10A (C) cells from different samples. B, D. Mean p53 target module score and HRAS log-normalized expression per condition in NDF (B) and MCF10A (D)cells. Bars are the mean, error bars the standard error of the mean. For each measure, the STP-minus-late-GR difference is shown above the bars with a 95 percent bootstrap interval over cells; the cell is the unit of analysis. STP was generated in a single independent experiment, so these are descriptive effect sizes, not replicate-level inference. E, F. Dynamic of expression of p53 target genes and HRAS. UMAP feature plots showing log-normalized expression of the indicated genes. Each point represents one cell; color indicates expression level. D, H. Presence of specific and alternative lineage signature markers in NDF (G) and MCF10A (H) cells. I, J. Expression of “Inflammation” or “response to Viruses” signatures in NDF (I) and MCF10A (J) cells. K. Mean module score for five programs: “Fibroblast Identity”, “Interferon Response”, “Inflammation”, HDAC targets”, and “p53 targets” — across both lineages; columns are EV, late-GR, and STP for NDF (left of the divider) and MCF10A (right); color scale, module score.

Thus, the transcriptional programs of STP fibroblasts continued to converge toward those of the original nontransformed state, whereas STP epithelial cells continued to diverge. The IFN response was the only program that showed the opposite trend and may contribute to higher fibroblast resistance to oncogenic transformation (23, 24).

## Discussion

A distinctive feature of this study is the integration of complementary approaches in a controlled human-cell system. Contrary to other studies exploring effects of oncogenes on p53 functional background, we disabled p53 from the moment of transduction. We followed oncogene-induced transformation longitudinally in mesenchymal and epithelial cells, rather than comparing only the starting and end states. Cellular barcoding allowed us to determine whether the observed changes reflected population-wide responses or selection of rare clones, while single-cell transcriptomics resolved the timing and heterogeneity of the associated molecular programs. We also excluded antibiotic selection to preserve cells with different response to oncogene transduction. Finally, the parallel analysis of dermal fibroblasts and mammary epithelial cells allowed us to distinguish shared responses to combined HRAS-G12V expression and p53 disruption from lineage-dependent transformation trajectories. Together, these features provide a framework for examining cell-intrinsic barriers to transformation independently of microenvironmental selection.

Our central observation is that oncogenic perturbation initiated a rapid, population-wide reprogramming response but did not drive all cells toward the same stable endpoint. Barcode complexity and entropy remained stable during the period of morphological transformation, with no evidence that a small number of clones expanded at the expense of the rest of the population. Consistent with this result, essentially all GR-transduced cells departed from the control transcriptional state. The earliest response in both lineages included loss of differentiation-specific markers, induction of RAS- and cancer-associated programs, inflammatory and interferon signaling and expression of alternative-lineage markers. This was accompanied by increased transciptional entropy, and chromatin decondensation. Many of these changes subsequently diminished. Some cells moved back toward the transcriptional state of the parental cells, whereas others continued to diverge or stabilized in distinct altered states. These findings argue against a model in which transformation is initiated exclusively in a rare preexisting permissive population. Instead, they support a model in which most cells can enter an oncogene-induced plastic state, but cell-intrinsic barriers influence whether that state resolves toward lineage recovery or progresses toward full transformation.

The observed reprogramming was not explained simply by changes in cell-cycle composition. In fibroblasts, control, early-GR, and late-GR populations remained transcriptionally distinct within both cycling and noncycling compartments, and the major module-score trajectories were concordant between these compartments. Chromatin accessibility also increased in both cycling and noncycling cells. Epithelial-cell profiles were influenced more strongly by cell-cycle state, and cell cycle and experimental batch accounted for more of their overall transcriptomic organization than they did in fibroblasts. Nevertheless, early GR-transduced epithelial cells also separated from their corresponding controls, and chromatin decondensation was not restricted to cycling cells. Thus, proliferation contributes to the structure of the single-cell data—particularly in epithelial cells—but does not account for the shared loss of lineage identity, inflammatory activation, increased entropy, or chromatin opening induced by GR.

Several features of the response were common to fibroblasts and epithelial cells. Both lineages entered an early state of reduced lineage fidelity, expressed markers normally associated with alternative differentiation programs, activated inflammatory and antiviral pathways, and showed transient increases in transcriptional entropy and chromatin accessibility. Both populations subsequently became heterogeneous, with some cells moving toward control-like profiles and others remaining distant from them. The acquisition of epithelial-like features by fibroblasts and mesenchymal features by epithelial cells can be viewed as parallel manifestations of reduced lineage constraint rather than unrelated lineage-specific phenomena. In both systems, therefore, HRAS activation and p53 disruption appeared to open a transient window of plasticity during which normally restricted transcriptional programs became accessible. The partial resolution of this state, without detectable loss of barcode diversity, suggests active adaptation within many clones rather than replacement of the population by a few resistant clones.

Despite these common features, the two lineages followed markedly different trajectories. Fibroblasts underwent larger and more discrete transcriptomic shifts that were evident independently of cell-cycle state. They initially lost fibroblast-identity programs and later separated into distinct states, including populations that partially restored lineage identity. After selection for a transformed phenotype, fibroblasts continued to move toward aspects of the original lineage program, showed lower HRAS expression, higher p53 targets expression less inflammation, but acquired a strong interferon-response signature. By contrast, epithelial cells changed more gradually and continuously. Selected epithelial populations became more separated from EV controls, showed further loss of epithelial identity, p53 targets, and reinforced HRAS, inflammatory, EMT, and TGF-β-associated programs. Thus, fibroblasts exhibited a more pronounced initial displacement but also stronger subsequent reversion, whereas epithelial cells underwent subtler, progressive changes that carried them farther from the normal epithelial state. Thus suggests the presence of a stronger lineage restricting barrier in cells of mesenchymal origin, than in epithelial cells.

The molecular basis of this barrier remains unknown, but our data suggest several, potentially connected mechanisms. First, the transient increase in single-cell entropy indicates that GR drives cells into a less committed state from which multiple outcomes are possible. Second, chromatin decondensation may provide a physical substrate for rapid changes in lineage, inflammatory, and alternative-differentiation programs. Loss of p53 could contribute because p53 participates in several mechanisms of chromatin silencing, whereas activated RAS has been linked to both chromatin compaction and decompaction in context-dependent settings (7, 8, 10, 25). Desilenced chromatin may also generate aberrant transcriptional products, including endogenous retroviral transcripts and double-stranded RNA, that engage pattern-recognition receptors and activate NF-κB and interferon-regulatory factors (26-28). Metabolic adaptation, proteotoxic stress, DNA-damage responses, and feedback attenuation of RAS signaling may provide additional safeguards. Importantly, chromatin opening could be either a driver or a consequence of transcriptional reprogramming. Dissecting its temporal and causal relationship to lineage loss and inflammatory signaling will therefore be essential.

The contrasting trajectories of fibroblasts and epithelial cells may also be relevant to the marked predominance of carcinomas over sarcomas among adult cancers (29). Our findings raise the possibility that mesenchymal cells can mount a stronger cell-intrinsic reversion program after oncogenic perturbation, whereas epithelial cells may be more likely to consolidate progressive lineage loss and transformation-associated programs. Such a difference could contribute to lineage-specific susceptibility to malignant transformation. However, cancer incidence is shaped by many additional variables, including tissue mass and turnover, exposure to mutagens, stem-cell organization, microenvironmental signals, immune surveillance, and the spectrum of initiating mutations. Our in vitro comparison of one fibroblast model and one mammary epithelial model cannot establish a causal connection between the observed trajectories and population-level cancer incidence. The sarcoma–carcinoma difference should therefore be regarded as a testable biological hypothesis generated by these data, rather than as an explanation established by the present study.

The interferon response is a particularly interesting candidate component of the transformation barrier. In selected fibroblast populations, restoration of fibroblast identity and reduced HRAS-associated activity coincided with a strong and persistent interferon-response program, whereas selected epithelial cells showed little interferon activation and continued to move away from the control state. This inverse association between interferon signaling and progression of the transformed transcriptional state is consistent with a protective role. Cell-intrinsic interferon signaling could restrict transformation through growth inhibition, senescence-like programs, innate immune surveillance, or elimination of cells producing aberrant nucleic acids (23, 24). Alternatively, the interferon signature may simply mark chromatin desilencing or cellular stress without contributing directly to resistance. Because the present data establishes correlation rather than causation, perturbation of interferon production, sensing, and downstream signaling will be required to determine whether this pathway actively promotes reversion or limits stable transformation.

Several limitations of our study should be considered. The experimental models were adapted to cell culture, and the epithelial models were immortalized; these features can alter senescence, apoptosis, lineage fidelity, stress tolerance, and baseline chromatin organization. Long-term culture may also enrich proliferative, stress-resistant cells before oncogene introduction. The GR construct models only one driver combination, and dominant-negative p53 may not reproduce all consequences of the diverse TP53 alterations found in tumors. Although TP53 and RAS-pathway alterations are among the most frequent oncogenic events, responses to other drivers, mutation orders, expression levels, and tissue contexts may differ. The lack of clonal selection during the observed interval does not exclude selection below the sensitivity of the barcode assay, selection occurring before the first measurement or during later passage, or nonproliferative forms of selection. Barcodes track heritable lineage, so the clonal data exclude selection of a stably distinct subpopulation. They cannot report a permissive state that individual cells enter and leave, since sister cells sharing a barcode need not share that state.

Some time-course comparisons were descriptive at the cell level, and selected transformed-phenotype populations were generated in a limited number of independent experiments. Finally, the in vitro design intentionally isolates cell-autonomous mechanisms and therefore does not capture extracellular matrix architecture, stromal interactions, immune surveillance, or other tissue-level constraints that influence tumor initiation in vivo.

Future studies should define the molecular components and causal order of the proposed barrier. Separating the effects of HRAS-G12V from those of p53 disruption, varying their order and dosage, and extending the analysis to additional driver combinations will reveal which responses are general features of transformation. Testing primary and organ-specific mesenchymal and epithelial cells from multiple donors will determine whether the lineage differences observed here are reproducible and whether they relate to sarcoma and carcinoma susceptibility. Lineage-resolved perturbation experiments targeting chromatin regulators, metabolic checkpoints, innate nucleic-acid sensors, and interferon signaling should establish which pathways control reversion or progression. Joint single-cell measurements of transcriptomes, chromatin accessibility, lineage barcodes, and spatial or phenotypic states would connect molecular trajectories directly to transformation outcomes. By distinguishing uniform early responses from lineage-dependent stable outcomes, the present framework offers a tractable route to identifying mechanisms that allow oncogene-bearing cells to remain nonmalignant and, potentially, to finding strategies that reinforce those mechanisms.

## Materials and Methods

### Details of all methods are provided in Supplementary Materials

Raw single-cell RNA-sequencing and clonal-barcode sequencing data generated in this study have been deposited in the Gene Expression Omnibus under accession no. GSE337416. Source data underlying the main and supplementary figures and custom Python analysis code are available from the corresponding author, Katerina V. Gurova (Roswell Park Comprehensive Cancer Center), upon request.

## Acknowledgments

We would like to express our gratitude to Cell Stress Biology administrative staff, Mary Morgan, Sarah Marcey, Vanessa Glover, Denise Pasinsky, Xin Chen and Bruce Specht for constant support of our research; our colleagues, Drs. Andrei Gudkov, Dominique Smiraglia, Eugene Kandel, Subhamoy Dasgupta and Veena Prahlad for critical discussion of the project. We acknowledge the RPCCC shared resources: Genomics, Bioinformatics, Biostatistics and Statistical Genomics, Gene Modulation Services, and Flow & Immune Analysis. This study was supported by funding from NIH/NCI R01CA266216 to KG. The shared resources of Roswell Park were supported by NCI CCSG P30CA016056, awarded to Roswell Park Comprehensive Cancer Center.

## Author Contributions

K.G. and C.F. designed research, performed experiments, analyzed data, wrote the paper; A.S., M.C., V.J., A.S., and B.B. performed research; P.S., J.L.: prepared NGS libraries and performed sequencing; K.L. provided reagents and protocols to use them; D.D., and J.W. helped with bioinformatic analysis;

