## Supplementary materials for "Oncogenic transformation proceeds through a transient state of cellular plasticity constrained by lineage-specific barriers"

**This PDF file includes:**

Supporting text
Figures S1 to S15
SI References

### Supporting Information Text

### Materials and Methods

#### Chemicals and Antibodies

Propidium iodide, hexadimethrine bromide (Polybrene), RNase A, and methylene blue were purchased from Sigma-Aldrich (St. Louis, MO). DAPI and the EU kit were purchased from Invitrogen/Thermo Fisher Scientific (Grand Island, NY). The DO-1 anti-p53 antibody was purchased from Santa Cruz Biotechnology (Santa Cruz, CA), and the Alexa Fluor 488–conjugated anti-mouse secondary antibody was purchased from Invitrogen/Thermo Fisher Scientific. CBL0137 was provided by Incuron, LLC.

#### Cell Lines and Culture Conditions

MCF10A cells were obtained from the American Type Culture Collection (ATCC). Primary normal human neonatal dermal fibroblasts (NDFs), pooled from three donors, were obtained from AllCells, LLC (Alameda, CA). NDFs were maintained in high-glucose DMEM (Invitrogen) supplemented with 5% fetal bovine serum (FBS) and antibiotics at 37 °C in a humidified atmosphere containing 5% CO2. MCF10A cells were grown in DMEM/F12 (Invitrogen #11330-032) with 5% Horse Serum (Invitrogen#16050-122), 20ng/ml of EGF (Peprotech, Rocky Hill, NJ), 0.5 mg/ml hydrocortisone (Sigma-Millipore, cat #H-0888), 100 ng/ml of Cholera Toxin (Sigma-Millipore, cat #C-8052), 10μg/ml of Insulin (Sigma-Millipore, cat #I-1882) and antibiotics.

#### Lentivirus Production and Transduction

Lentiviral particles were produced by the Roswell Park Gene Modulation Shared Resource by transfecting HEK293 cells with the packaging plasmid psPAX2 (encoding gag, pol, and rev), the VSV-G envelope plasmid pMD2.G, and either a lentiviral vector encoding the dominant-negative p53 mutant GSE56 and the HRAS-G12V oncogene separated by an internal ribosome entry site (IRES) [28] or the corresponding empty-vector (EV) control. Viral supernatants were titrated separately in NDF and MCF10A cells to determine the dose corresponding to approximately one transducing unit (TU) per cell. Transduced cells were identified 72 h after transduction by immunostaining for p53 followed by flow cytometry or quantitative imaging, as described below.

#### Cell-Proliferation and Transformation Assays

For two-dimensional (2D) growth assays, 50,000 cells were plated per 30-mm dish or per well of a six-well plate in triplicate and cultured for 3 days. Cells were stained with methylene blue and photographed. For quantification, the dye was solubilized in 1% SDS, and absorbance at 600 nm was measured with a microplate reader.

For colony-formation assays, 100,000 NDFs were plated per 100-mm dish and cultured for 7 to 10 d, until colonies of GR-transduced cells became visible. Plates were then stained with methylene blue and photographed. For growth in semisolid medium, 100,000 cells were suspended in 0.3% agarose prepared in complete culture medium and overlaid onto a solidified layer of 0.6% agarose in six-well plates. After 1 h, 1 mL of culture medium was added above the top agarose layer. Colonies were counted and photographed after 2 wk. To return cells to 2D culture, the top agarose layer containing colonies was collected, diluted with prewarmed medium, and transferred to standard six-well plates without a bottom agarose layer. Attached cells were photographed after 24 h.

**Immunofluorescence Staining**

Transduced and untransduced cells were stained with the DO-1 anti-p53 primary antibody and an Alexa Fluor 488–conjugated anti-mouse secondary antibody using a manufacturer protocols. Cells were stained in suspension for flow cytometry and in 96-well plates for quantitative imaging (Greiner Bio-One, Monroe, NC; catalog no. 655090). Untransduced cells treated with 1 µM CBL0137 were used as positive controls for p53 staining.

**Chromatin-Accessibility Imaging**

At the indicated times after transduction, 3,000 cells were plated per well in black, clear-bottom 96-well plates (Greiner Bio-One; catalog no. 655090). The following day, the medium was removed, and cells were fixed and permeabilized for 10 min at room temperature with 4% paraformaldehyde (PFA) in PBS containing 0.1% Triton X-100. Wells were washed once with PBS, filled with PBS, sealed, and stored at 4 °C until all time points had been collected. Cells were then stained overnight at room temperature with RNase A (100 µg/mL) and propidium iodide (1 µg/mL).

**Microscopy and Image Analysis**

Phase-contrast images were acquired with a Zeiss Axio Observer A1 inverted microscope equipped with a Zeiss MRC5 camera and AxioVision Rel. 4.8 software. Quantitative images were acquired with a Cytation 5 automated imager (BioTek/Agilent Technologies, Santa Clara, CA) using a 4× objective and per-well autofocus. Each well image was assembled from four fields. Images were processed and analyzed with Gen5 Image Prime software (BioTek/Agilent). Approximately 300 to 5,000 objects were analyzed per well. Object masks were restricted to nuclei. Before analysis, objects from all wells were sorted by size; objects smaller than 15 µm or larger than 40 µm were excluded to remove debris and cell clusters.

**Lentiviral Cellular Barcoding and Sequencing**

Custom CloneTracker lentiviral libraries containing approximately 1 million unique 30-nt barcodes, mCherry, and a puromycin-resistance gene were provided by Cellecta (Mountain View, CA). Cells were transduced at 0.3 TU per cell, as determined from the proportion of mCherry-positive cells, selected with 2 μg/mL puromycin for 48 h, expanded, and cryopreserved. The same frozen cell stock was used for each experiment. To assess population complexity, $1 \times 10^6$ cells were collected at the indicated time points and snap-frozen as pellets. Genomic DNA was isolated with the DNeasy Blood & Tissue Kit (QIAGEN, Germantown, MD; catalog no. 69504). Sequencing libraries were prepared by a two-step PCR procedure targeting the lentiviral barcode sequences. Primers complementary to the constant regions flanking the barcode were appended with the forward adapter sequence TCGTCGGCAGCGTCAGATGTGTATAAGAGACAG or reverse adapter sequence GTCTCGTGGGCTCGGAGATGTGTATAAGAGACAG required for the second PCR.

For the first PCR, the maximum available amount of genomic DNA (up to 50 μg) was used to amplify each target region. Amplicons were purified with AMPure XP beads (Beckman Coulter, Brea, CA) and analyzed with a TapeStation DNA 1000 system (Agilent, Santa Clara, CA). First-round amplicons were subjected to eight additional PCR cycles with the Nextera Index Kit (Illumina, San Diego, CA). These primers anneal to the adapter overhangs incorporated during the first PCR and add a unique index to each sample, enabling library pooling and multiplexed sequencing. Indexed libraries were purified with AMPure XP beads, quantified by KAPA qPCR (KAPA Biosystems, Wilmington, MA), and pooled in equimolar amounts. Libraries were sequenced with 100-cycle paired-end reads on an Illumina NovaSeq 6000 at the Genomics Shared Resource, Roswell Park Comprehensive Cancer Center, according to the manufacturer's protocol.

**Barcode Quantification and Clonal-Diversity Analysis**

Sequencing reads were processed with custom software that assigned reads to CloneTracker barcode references and generated a barcode-count matrix for each sample. Raw counts were used for all diversity calculations, and a barcode was considered present when supported by at least 10 reads. To compare samples sequenced at different depths, each sample was rarefied by multinomial subsampling to the minimum depth within its lineage (1.5 million reads for NDF and 3.8 million reads for MCF10A). Clonal diversity was quantified as the Shannon entropy of the barcode-frequency distribution, $H = -\Sigma_i p_i \ln(p_i)$, using scipy.stats.entropy in SciPy (1). Ninety-five percent confidence intervals were estimated from 10,000 bootstrap resamples drawn with replacement. EV and GR populations were compared at each time point using two-sided Mann–Whitney U tests, with Benjamini–Hochberg correction across comparisons within each lineage. The analysis followed the approach of Bhang et al (2).

**Single-Cell RNA Sequencing and Analysis**

Cells were plated at 1 × $10^6$ cells per 150-mm dish. The following day, cells were transduced with GR lentivirus at 1 TU per cell or with an equivalent amount of EV lentivirus in the presence of 10 μg/mL Polybrene. After 24 h, the medium was replaced with virus-free medium. Every 3 to 4 d, when cultures approached confluence, cells were trypsinized and counted. For each time point, 4 × $10^6$ cells were fixed with the Evercode Cell Fixation Kit (Parse Biosciences, Seattle, WA) according to the manufacturer's protocol; the remaining cells were replated at 1 × $10^6$ cells per 150-mm dish. Fixed single-cell suspensions were stored at −80 °C until all samples from an experiment had been collected. Libraries were prepared by split-pool combinatorial barcoding with the Parse Biosciences Evercode Whole Transcriptome kit. Approximately 100,000 cells representing 12 samples were sequenced per experiment. Two independent experiments were performed with NDFs and three with MCF10A cells. Libraries were sequenced on an Illumina NovaSeq 6000 at the Genomics Shared Resource, Roswell Park Comprehensive Cancer Center, to a mean depth of approximately 30,000 reads per cell.

FASTQ files were generated from Illumina base-call files with bcl2fastq v2.20. Alignment, sample annotation, read counting, and count-matrix generation were performed with the Parse Biosciences split-pipe pipeline v0.9.6p. Quantification, normalization, and visualization were performed with Scanpy {Wolf, 2018 #2267}, Seaborn, and Matplotlib. Statistical analyses were performed with SciPy {Virtanen, 2020 #2268}.

Cells for which mitochondrial-gene transcripts accounted for more than 12% of total counts were excluded. Counts were normalized to 10,000 counts per cell and log-transformed. Highly variable genes were selected with the following parameters: min_mean = 0.0125, max_mean = 3, and min_dispersion = 0.25. Principal-component analysis was performed on the selected genes. A neighborhood graph was constructed from the first 30 principal components using 10 neighbors per cell, embedded with uniform manifold approximation and projection (UMAP), and partitioned with the Leiden algorithm at a resolution of 0.5. Cluster-marker genes were identified with the Wilcoxon rank-sum test. Per-cell module scores for curated gene sets—including fibroblast identity, inflammation, interferon response, HDAC1/2/3 targets, p53 targets, additional cancer-associated programs, and S- and G2/M-phase cell-cycle signatures—were calculated with the Scanpy score_genes function. Over-representation of MSigDB C2 gene sets (v2024.1) among cluster- and time point–specific marker lists was assessed with the Enrichr method implemented in GSEApy {Fang, 2023 #2266}. All single-cell analyses were performed in Python with Scanpy.

#### Single-Cell Transcriptome-Entropy Analysis

Single-cell network entropy was calculated for each cell with the Correlation of Connectome and Transcriptome (CCAT) statistic (Teschendorff and Enver, REF), defined as the Pearson correlation between log-normalized gene expression and protein–protein interaction connectivity. A Python implementation, validated against the reference R routine to machine precision, was applied to a high-confidence, experimentally supported subset of the STRING v12 network (Szklarczyk et al., REF). Counts were normalized to 10,000 counts per cell and log-transformed, and cells were processed in blocks of 5,000. As a network-independent measure of transcriptional plasticity, per-cell Shannon entropy was calculated from the normalized count distribution across detected genes. Because gene-detection rates differed among sequencing runs, GR and EV cells were compared only within the same experiment. Comparisons used two-sided Mann–Whitney U tests with Benjamini–Hochberg correction and 95% confidence intervals estimated from 10,000 bootstrap resamples.

#### Statistical Analysis and Figure Preparation

All experiments were performed at least twice and included at least two replicate wells. The value for each well was treated as a technical replicate and used to calculate the mean across replicate wells. For object-level analysis, objects from replicate wells were pooled. Differences between conditions were evaluated by unpaired t tests implemented with scipy.stats.ttest_ind in SciPy [29].

All computational analyses were performed in Python. Single-cell metrics and clonal-diversity measurements were compared using two-sided Mann–Whitney U tests, with Benjamini–Hochberg correction applied within each analysis. Bootstrap 95% confidence intervals were calculated from 10,000 resamples for clonal-diversity and entropy comparisons and from 5,000 resamples for module-score time courses. A fixed random seed of 42 was used for all sampling procedures. Plots were generated with Matplotlib and Seaborn.

#### Data, Materials, and Software Availability

Raw single-cell RNA-sequencing and clonal-barcode sequencing data generated in this study have been deposited in the Gene Expression Omnibus under accession no. GSE337416. Source data underlying the main and supplementary figures and custom Python analysis code are available from the corresponding author, Katerina V. Gurova (Roswell Park Comprehensive Cancer Center;), upon request.

**Supplementary Figures**

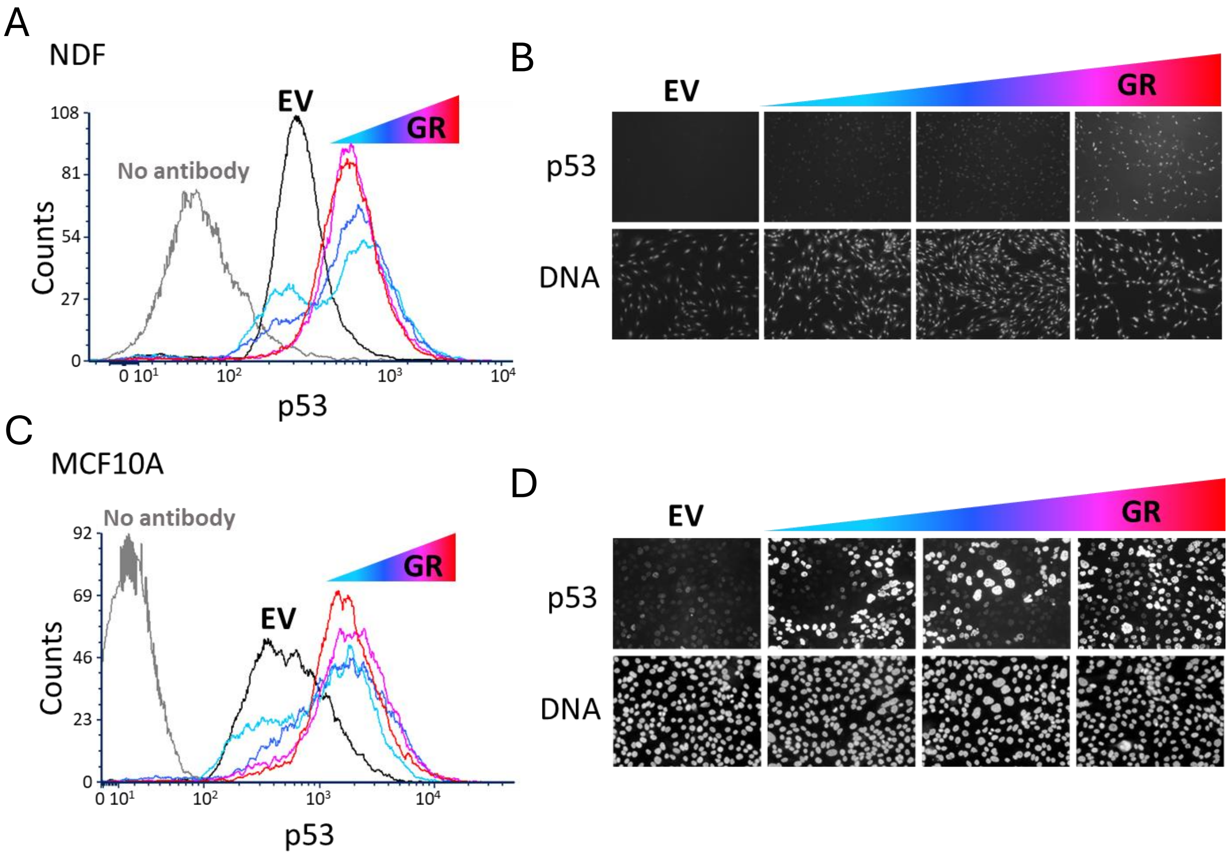


**Supplementary Figure S1.** Monitoring of transduction efficiency via staining of p53. NDF (A, B) and MCF10A (C, D) cells were transduced with different doses of GR or EV viruses. 72 hours later, cells were collected and stained with p53 antibody. A, C. Histograms of p53-related fluorescence acquired using flow cytometry. Grey line – unstained cells, black – EV transduced cells, colored lines – increasing concentrations of GR virus. Light and dark blue lines correspond to samples with some proportions of p53 negative cells, whose fluorescence was not different from EV cells, while purple and red samples have no such cells. The dose of GR virus corresponding to the purple line was used in further experiments. B, D. Immunofluorescence images of the same cells stained in plates. Cells were counterstained with Hoechst 33342 (DNA).

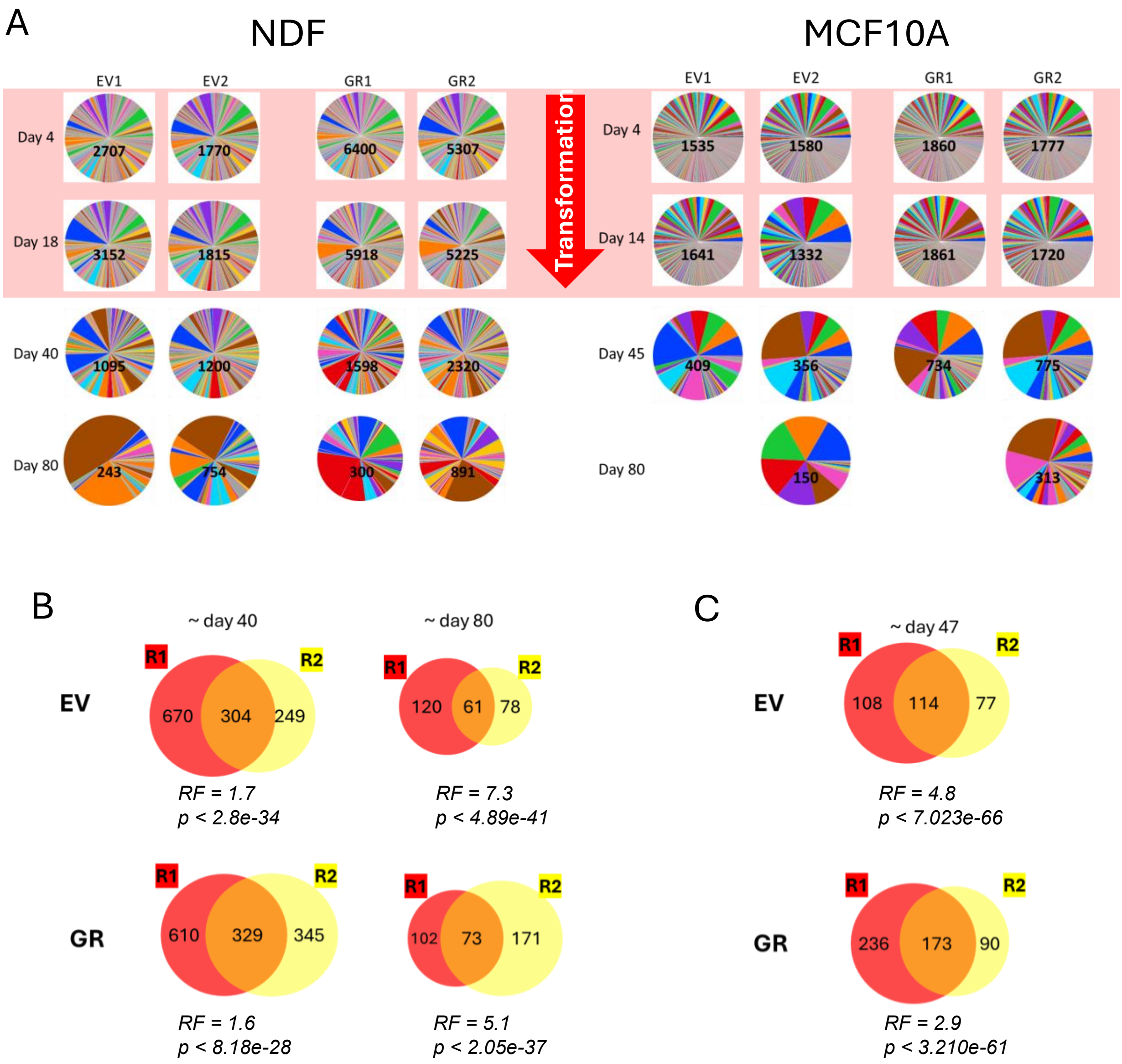


**Supplementary Figure S2.** Changes in population complexity of NDF and MCF10A cells after EV or GR transduction, assessed via barcode sequencing. A. Pie charts showing the abundance of random barcodes in cell populations at different time points after transduction. The results of two independent experiments are shown. The number on the pie chart is the total number of barcodes detected. The pink shade highlights the period of oncogenic transformation. B, C. Overlap in the barcodes between replicate experiments (R1 and R2) for GR and EV NDF (B) or MCF10A (C) cells at different time points after transduction. RF - Representation Factor, all p-values are significant with no difference between EV and GR cells.

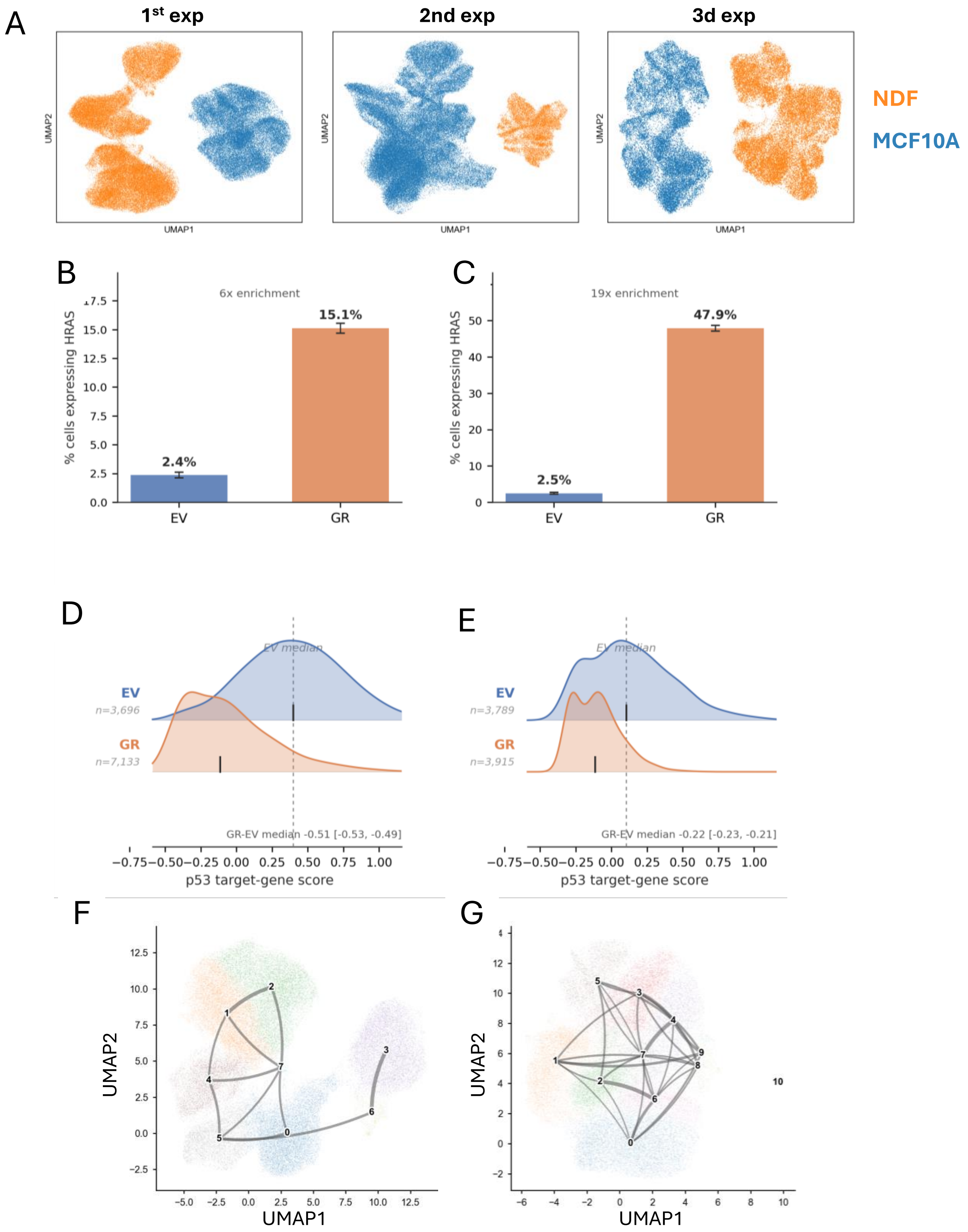


**Supplementary Figure S3. A.** Uniform manifold approximation and projection (UMAP) of all sequenced cells (n = 68,657 NDF; n = 113,720 MCF10A) in three independent experiments colored by lineage. B, C. Proportion of cells with detected HRAS corresponding reads in NDF (B)

and MCF10A (C) cells. D, E. Ridge plot of log-normalized expression of p53 target genes per cell in NDF (D) and MCF10A (E) cells st day 4 after transduction. Curves show kernel density estimates; vertical bar at the top of each ridge marks the EV-Day-0 median for reference. Genes are 10 canonical p53 targets: CDKN1A, MDM2, BBC3, BAX, GADD45A, TP53I3, PMAIP1, TIGAR, SESN1, SESN2.

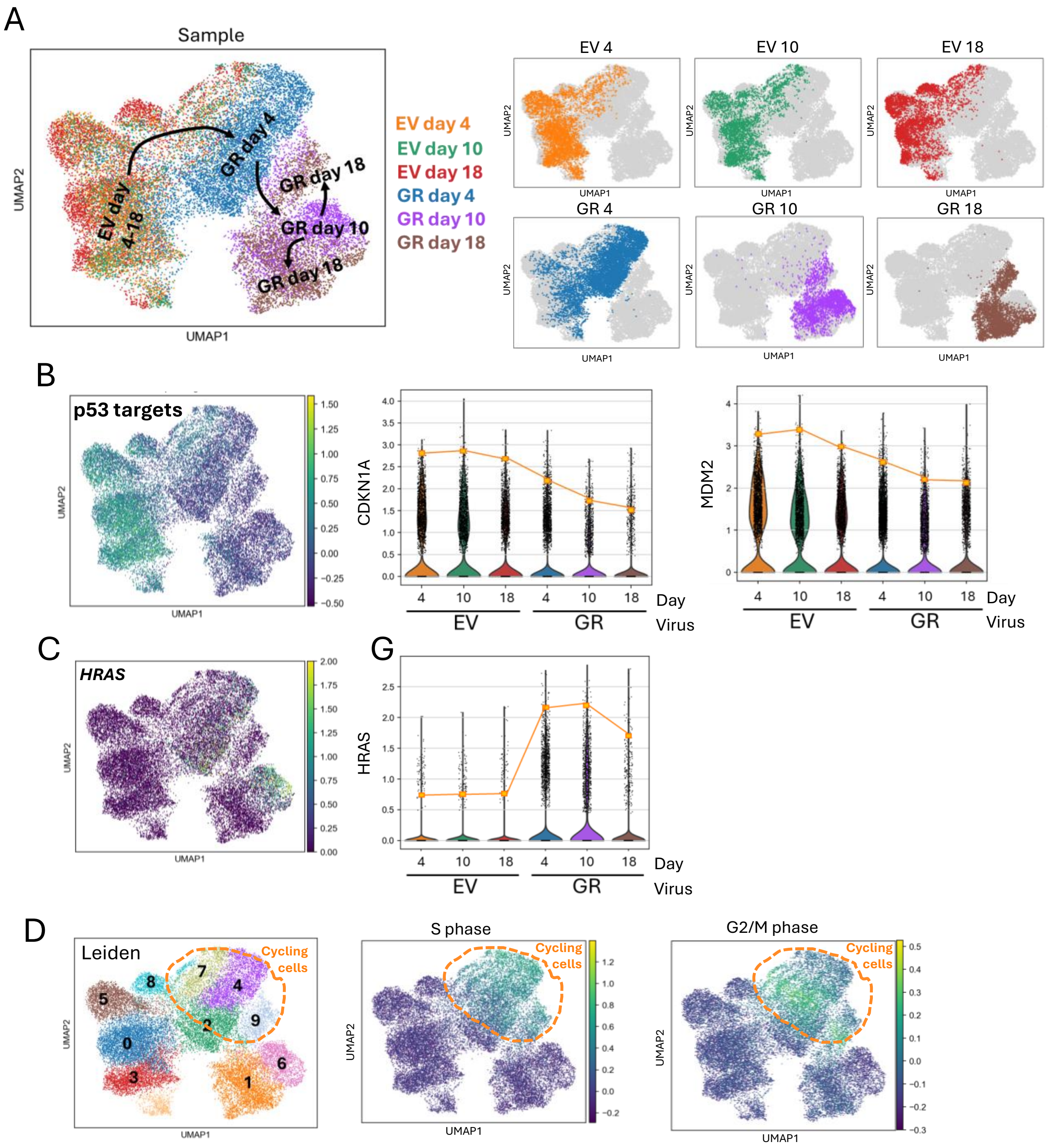


**Supplementary Figure S4.** Analysis of scRNA-seq of NDF cells in the third experiment. A. Distribution of all samples on UMAP plot (left) and position of individual samples against all others, grey dots (right). Number after EV or GR indicates the day of sample collection after transduction. B. Expression of p53 target genes on UMAP plot (left) and violin plots with two individual genes, CDKN1A (center) or MDM2 (right). C. Distribution of HRAS expression on UMAP and violin plots. D. UMAP plots with Leiden (left) clusters of NDF cells (left) and the presence of gene markers of S (central) or G2/M (right) phases of cell cycle. Orange square on all violin plots represents the value below which 90% of a cells with corresponding gene expression fall. Orange line shows tendency of corresponding gene expression in population of cells in each sample. Day 0 corresponds to control EV transduced cells. Orange dashed circles on D show cycling cells.

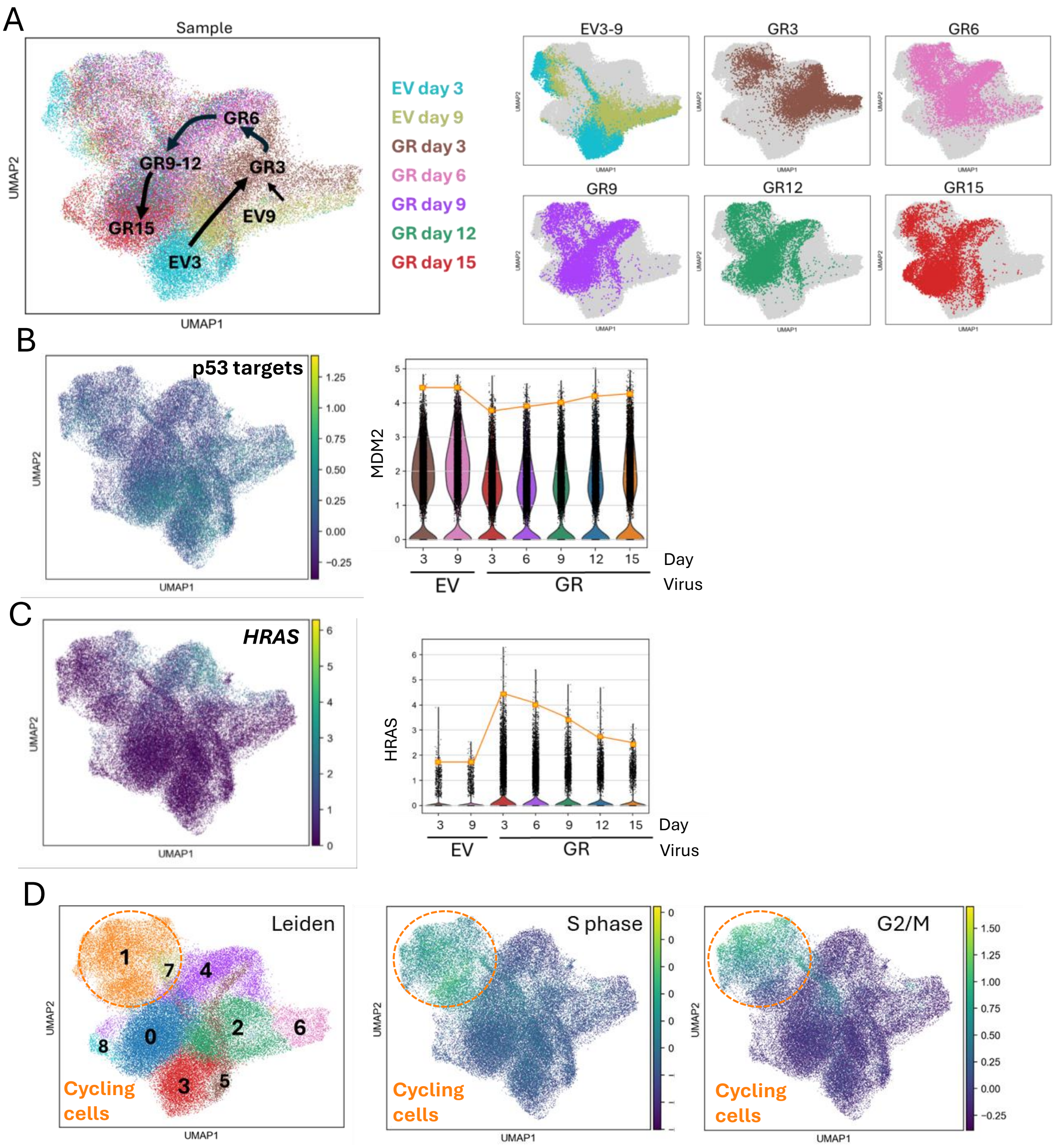


**Supplementary Figure S5.** Analysis of scRNA-seq of MCF10A cells in the second experiment. A. Distribution of all samples on UMAP plot (left) and position of individual samples against all others, grey dots (right). Number after EV or GR indicates the day of sample collection after transduction. B. Expression of p53 target genes on UMAP plot (left) and violin plots of MDM2 (right). C. Distribution of HRAS expression on UMAP and violin plots. D. UMAP plots with Leiden (left) clusters of MCF10A cells (left) and the presence of gene markers of S (central) or G2/M (right) phases of cell cycle. Orange square on all violin plots represents the value below which 90% of a cells with corresponding gene expression fall. Orange line shows tendency of corresponding gene expression in population of cells in each sample. Orange dashed circles on D show cycling cells.

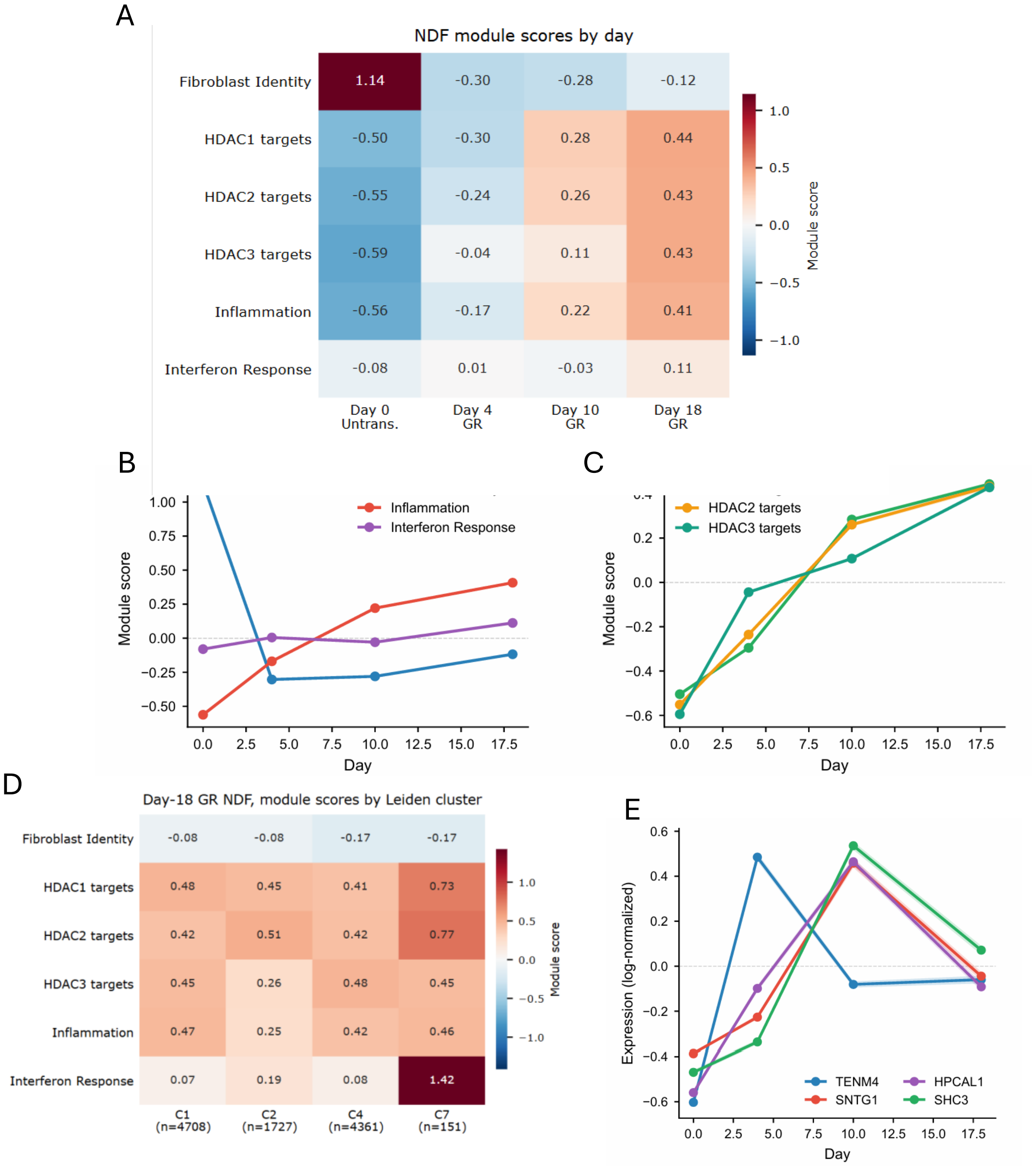


**Supplementary Figure S6.** NDF fibroblasts undergo coordinated reprogramming of fibroblast identity, chromatin remodeling, and inflammation during GR transformation. A. Heatmap of curated module scores (rows) across NDF cells stratified by day post-transduction (columns: EV Day 0, then GR Days 4, 10, 18). Scores were computed using sc.tl.score_genes (Scanpy v1.10) on log-normalized expression and z-scored across columns. B. Module dynamics over the time course, plotted as median module score per day with bootstrap 95% confidence intervals (CIs) as shaded ribbons (n = 2,000 resamples per timepoint, fixed random seed 42). C. HDAC1, HDAC2, and HDAC3 target module scores plotted as in panel B. D. Module scores stratified by Leiden cluster among Day-18 GR cells, shown as median ± interquartile range across cells per cluster. E. Neuronal-lineage gene expression (TENM4, SNTG1, HPCAL, SHC3) over the time course, plotted as median log-normalized expression with bootstrap 95% CIs.

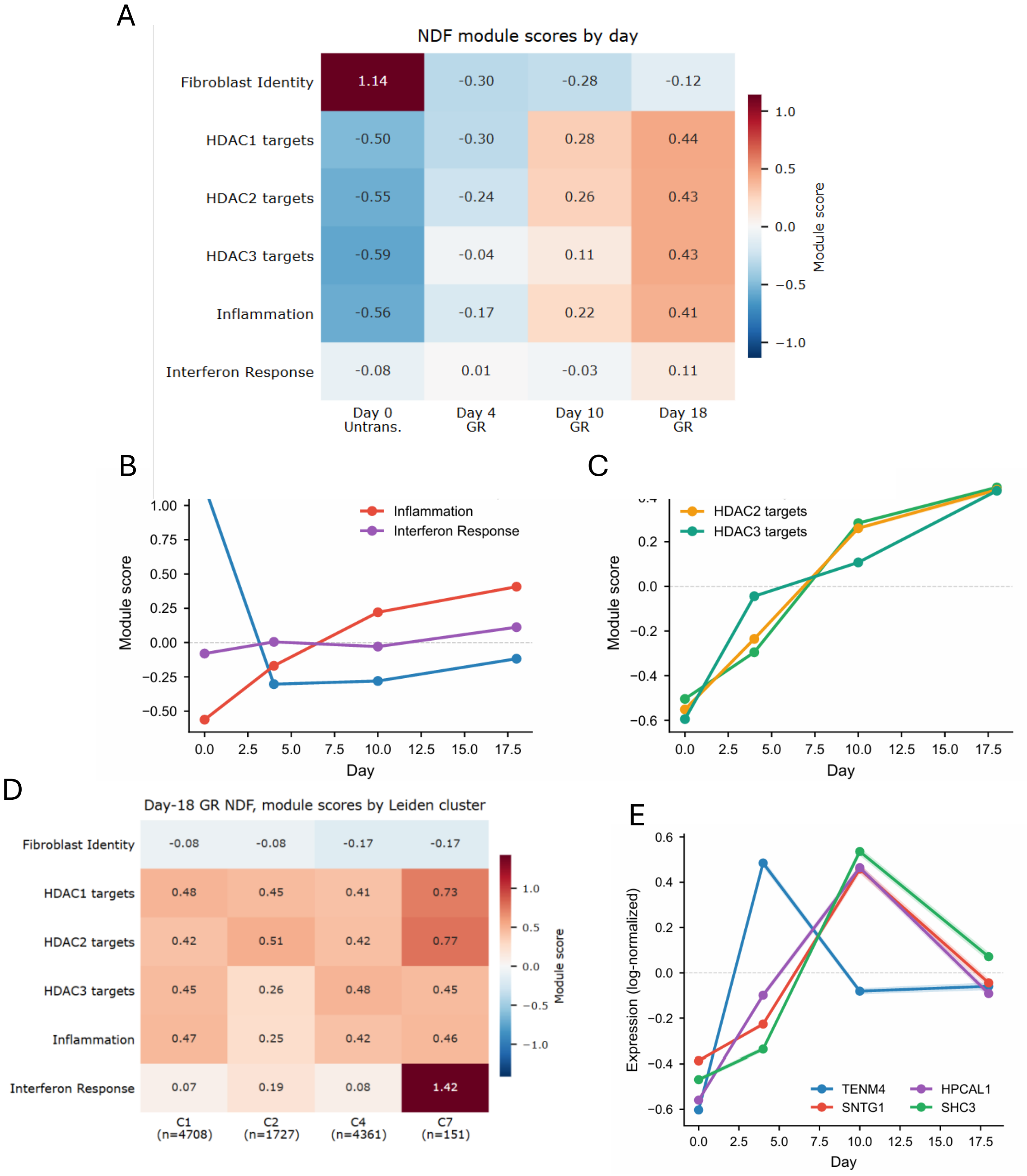


**Supplementary Figure S6.** NDF fibroblasts undergo coordinated reprogramming of fibroblast identity, chromatin remodeling, and inflammation during GR transformation. A. Heatmap of curated module scores (rows) across NDF cells stratified by day post-transduction (columns: EV Day 0, then GR Days 4, 10, 18). Scores were computed using sc.tl.score_genes (Scanpy v1.10) on log-normalized expression and z-scored across columns. B. Module dynamics over the time course, plotted as median module score per day with bootstrap 95% confidence intervals (CIs) as shaded ribbons (n = 2,000 resamples per timepoint, fixed random seed 42). C. HDAC1, HDAC2, and HDAC3 target module scores plotted as in panel B. D. Module scores stratified by Leiden cluster among Day-18 GR cells, shown as median ± interquartile range across cells per cluster. E. Neuronal-lineage gene expression (TENM4, SNTG1, HPCAL, SHC3) over the time course, plotted as median log-normalized expression with bootstrap 95% CIs.

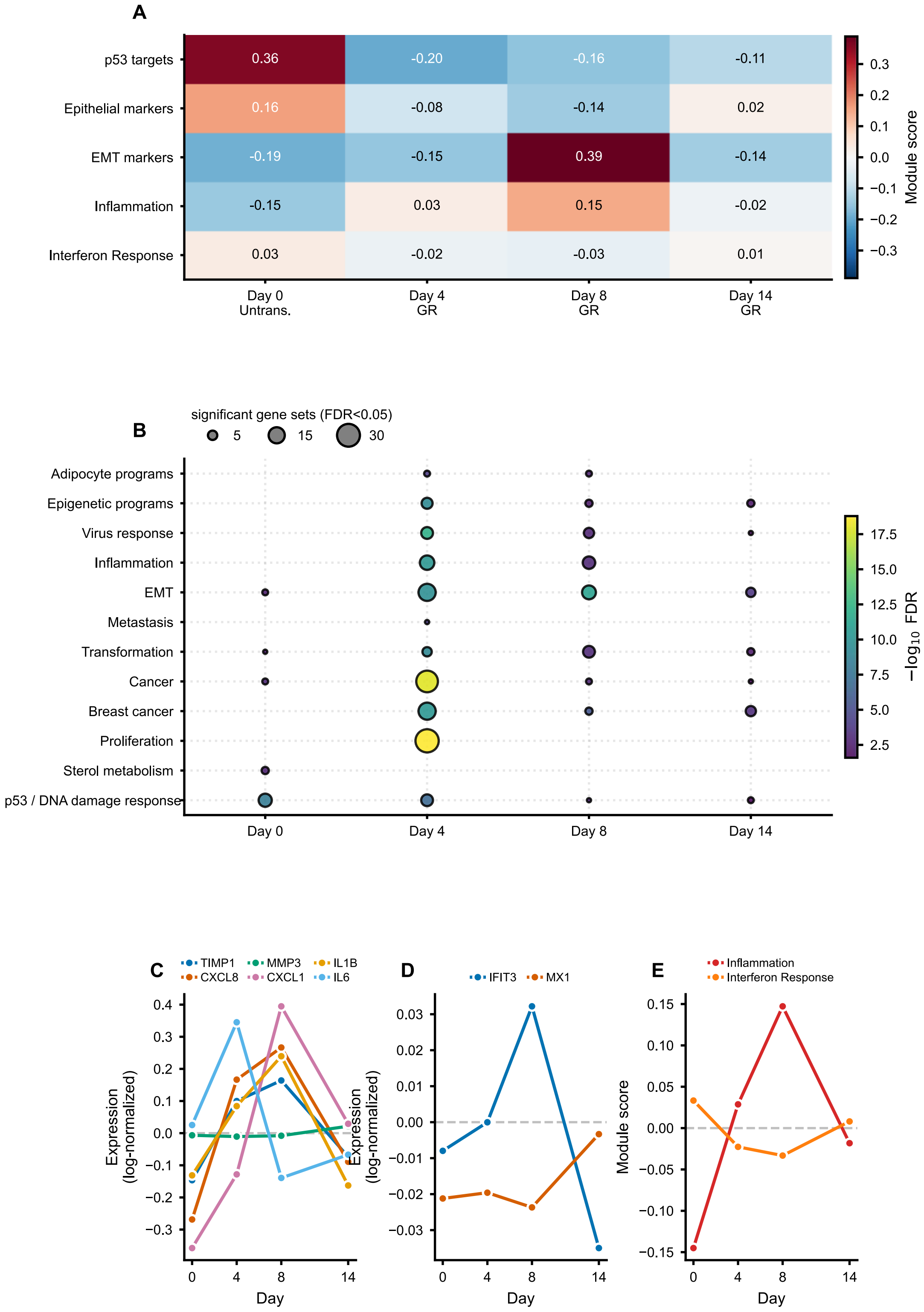


**Supplementary Figure S7.** Reprogramming of epithelial gene transcription during GR transformation. A. Enrichment heatmap of MCF10A programs across the time course. B.

Classification by MCF10A cluster (ORA; Enrichr, MSigDB C2). C. Individual NF-κB-regulated inflammatory genes (TIMP1, CXCL8, MMP3, CXCL1, IL1B, IL6). D. Individual IFN-responsive genes (IFIT3, MX1, ISG15) with distinct kinetics from the NF-κB-regulated set. Panel B: dot size, number of enriched gene sets at FDR < 0.05; color, maximum minus-log10 FDR; ORA (Enrichr, MSigDB C2). Panels C to E: one library per day; effects shown descriptively without significance testing (no error bars).

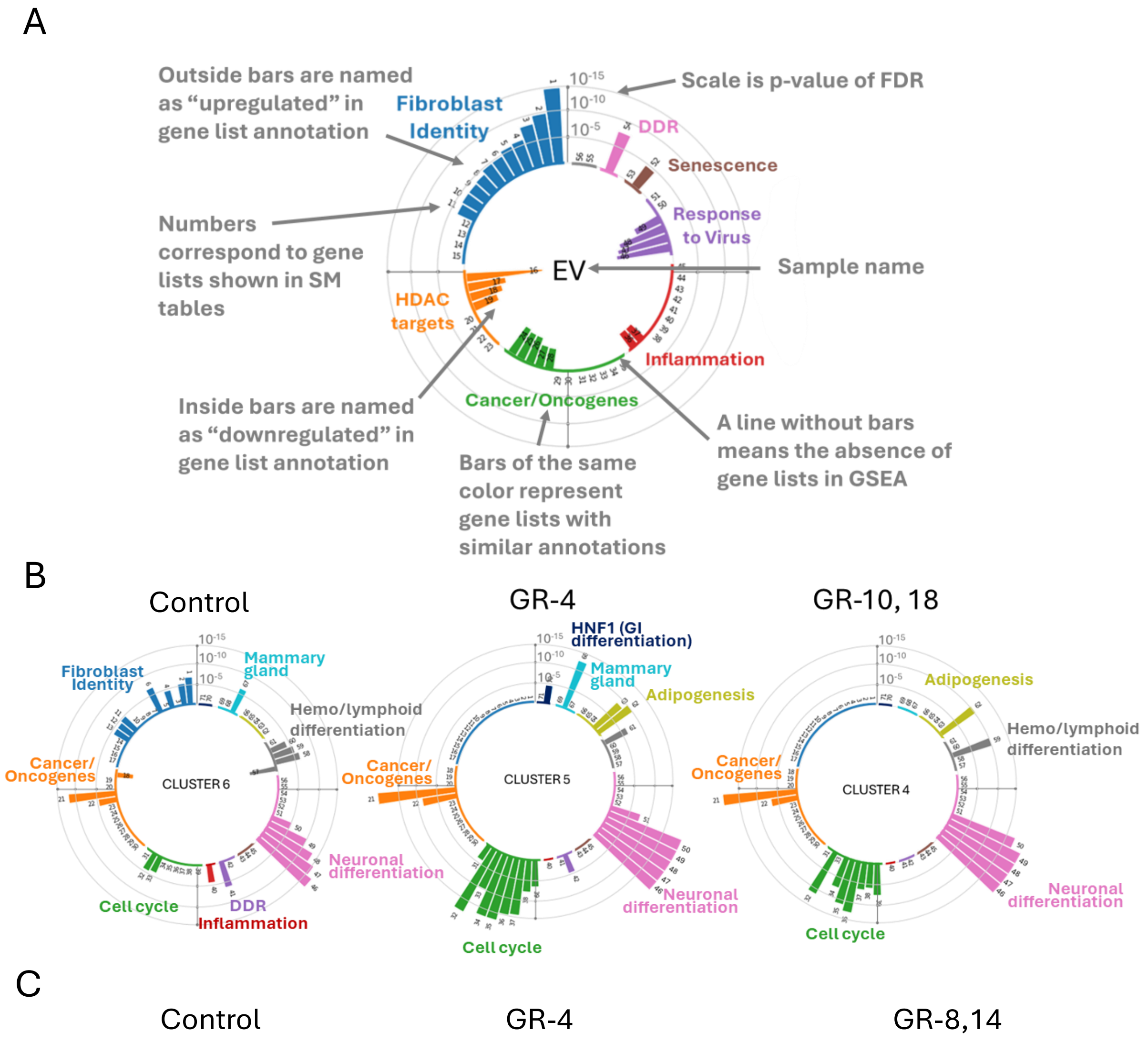


**Supplementary Figure S8.** A. Explanation of circular barplot components. B. Circular barplots of cycling NDF cells. C. Circular barplots of cycling MCF10A cells. Cluster number is shown in the center of circle. Numbers after GR – days of sample collection. Some clusters contain cells of different samples.

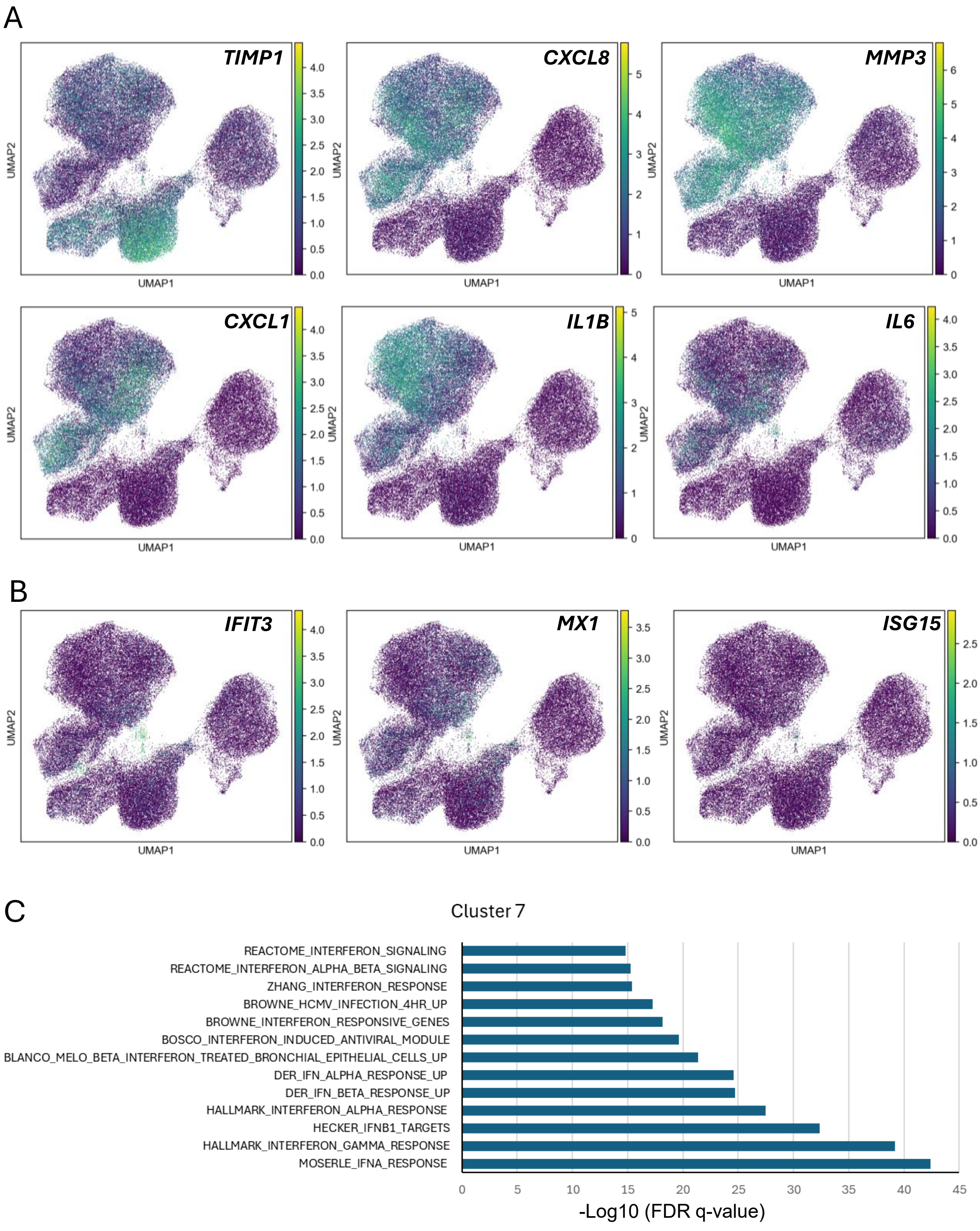


**Supplementary Figure S9.** Expression of inflammatory and interferon responsive genes in NDF cells. A. UMAP feature plots showing log-normalized expression of the indicated genes, targets of NF-kappaB. Each point represents one cell; color indicates expression level. N. The same plots for interferon responsive genes. C. GSEA of gene markers of Leiden cluster 7.

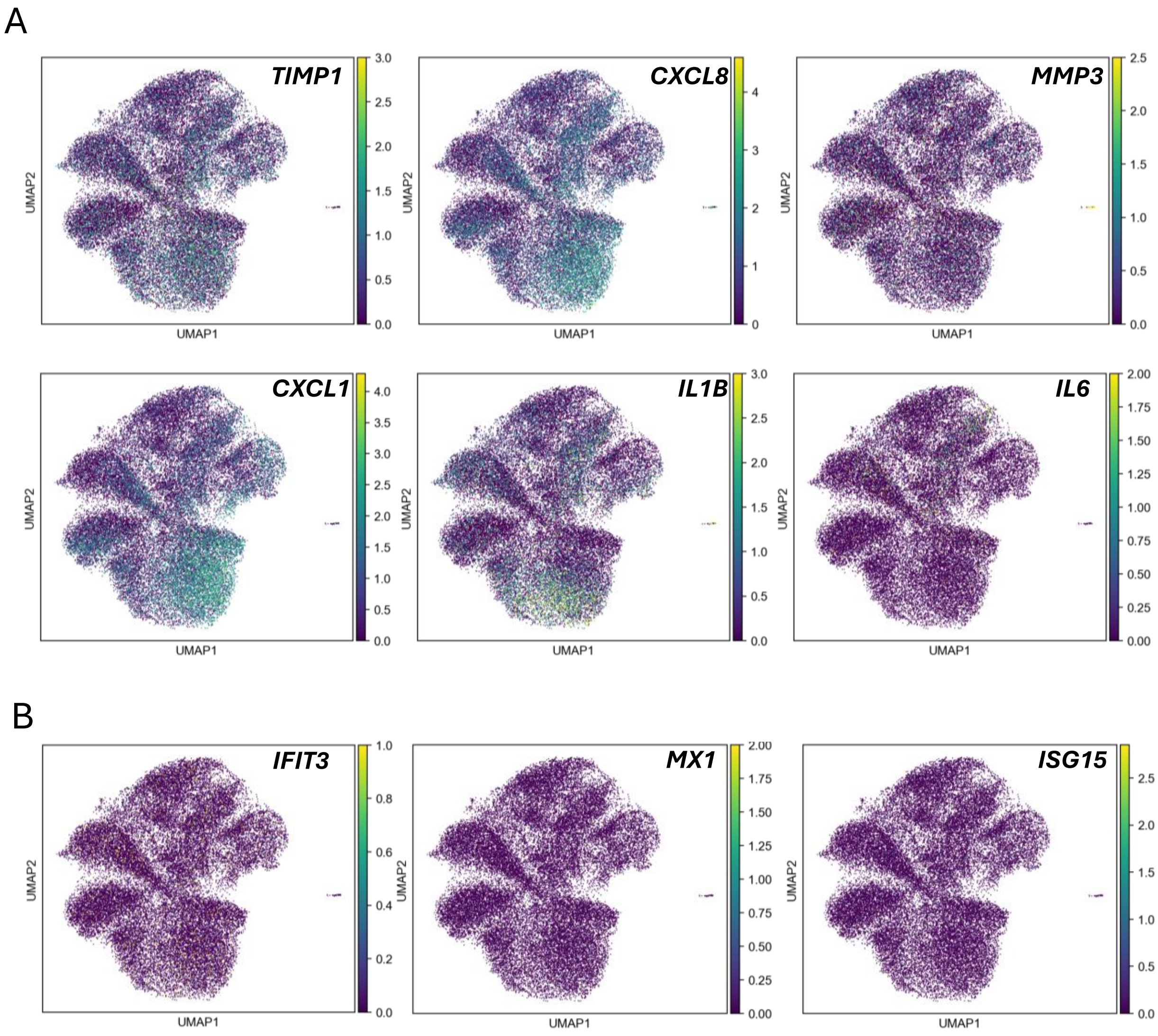


**Supplementary Figure S10.** Expression of inflammatory and interferon responsive genes in MCF10A cells. A. UMAP feature plots showing log-normalized expression of the indicated genes, targets of NF-kappaB. Each point represents one cell; color indicates expression level. N. The same plots for interferon responsive genes.

### Tables

**Table S1**. Sample inventory for single-cell RNA-sequencing experiments. Complete list of NDF and MCF10A samples profiled by SPLiT-seq, including condition (EV or GR), time point post-transduction (in days), replicate experiment number, and cell counts per condition after quality filtering. NDF experiments were performed in duplicate and MCF10A experiments were performed in triplicate. See Materials and Methods for cell preparation and library construction details.

**Table S2**. Lists of gene sets enriched in different groups of cells (samples and Leiden clusters) in order used for building circular plots with numbers corresponding to numbers on circular plots. GSEA analysis of the first experiment. File contains four sheets:
Samples NDF_1 – gene sets enriched in NDF samples from different days, corresponds to Fig. 4A.
Leiden Clusters NDF_1 – gene sets enriched in Leiden clusters of NDF cells, corresponds to Fig. 4E and S8B.
Samples MCF10A_1 - gene sets enriched in MCF10A samples from different days, corresponds to Fig. 5A.
Leiden Clusters_MCF10A_1 - gene sets enriched in Leiden clusters of MCF10A cells, corresponds to Fig. 5D and S8C.
