## Supplementary Table S1 for "Oncogenic transformation proceeds through a transient state of cellular plasticity constrained by lineage-specific barriers"

**Table S1. Sample design and single-cell RNA-sequencing summary for HRAS-G12V-induced transformation in NDF fibroblast and MCF10A epithelial cells.**

| **Lineage** | **Sample ID** | **Experiment** | **Day** | **Condition** | **Cells (n)** |
| --- | --- | --- | --- | --- | --- |
| NDF | NDF_Cont | First | 0 | EV | 9,996 |
| NDF | NDF_4d | First | 4 | GR | 16,762 |
| NDF | NDF_10d | First | 10 | GR | 17,795 |
| NDF | NDF_18d | First | 18 | GR | 11,103 |
| NDF | NDF-GRE-Hras colonies | Second | 50 | STP | 5,413 |
| NDF | NDF-GRE-Hras 35 days | Second | 50 | STP | 7,588 |
| **NDF subtotal** |  |  |  |  | **68,657** |
| MCF10A | MCF10A_Cont | First | 0 | EV | 9,895 |
| MCF10A | MCF10A-EV day3 | Second | 3 | EV | 8,402 |
| MCF10A | MCF10A GRE day3 | Second | 3 | GR | 8,207 |
| MCF10A | MCF10A_4d | First | 4 | GR | 5,151 |
| MCF10A | MCF10A GRE day6 | Second | 6 | GR | 7,919 |
| MCF10A | MCF10A_8d | First | 8 | GR | 9,120 |
| MCF10A | MCF10A-EV day9 | Second | 9 | EV | 8,543 |
| MCF10A | MCF10A GRE day9 | Second | 9 | GR | 7,058 |
| MCF10A | MCF10A GR day12 | Second | 12 | GR | 7,954 |
| MCF10A | MCF10A_14d | First | 14 | GR | 9,181 |
| MCF10A | MCF10A GR day15 | Second | 15 | GR | 9,092 |
| MCF10A | MCF10A 5% DM GR | Second | 50 | STP | 7,716 |
| MCF10A | MCF10A GR 35 days | Second | 50 | STP | 8,157 |
| MCF10A | MCF10A-GR colonies | Second | 50 | STP | 7,325 |
| **MCF10A subtotal** |  |  |  |  | **113,720** |
| **Total** |  |  |  |  | **182,377** |

**Condition codes.** EV: empty vector control (uninfected baseline at Day 0; vector-transduced for paired comparisons at later time points). GR: HRAS-G12V transduced cells progressing through the transformation time course. STP: cells selected for the transformed phenotype at Day 50 by anchorage-independent colony formation (GR-3D, sample IDs containing 'colonies'), growth-factor-independent proliferation in two-dimensional culture (GR-2D, MCF10A only; sample 'MCF10A 5% DM GR'), or continuous standard culture for 35 days (GR-LP, sample IDs containing '35 days').

**Experiment codes.** Each unique sample derives from one of two independent transduction experiments performed on different dates with separately thawed cell stocks. **First** denotes the primary time-course experiment for each lineage. **Second** denotes the experiment that included the Day 50 STP samples and additional intermediate time points for MCF10A.

**Sequencing platform and library preparation.** All samples were profiled by Parse Biosciences SPLiT-seq using three rounds of combinatorial barcoding (obs columns bc1_well, bc2_well, bc3_well). Libraries were sequenced on the Illumina NovaSeq platform with paired-end reads. Reads were aligned to the GRCh38 (hg38) human reference genome and quantified with the Parse Biosciences split-pipe pipeline (REF).

**Quality control thresholds.** Cells were retained if they passed all of: gene count ≥ 200, transcript count ≥ 500, mitochondrial transcript fraction ≤ 25%, and doublet detection (Scrublet) score below the cutoff selected per sample. After filtering, the NDF dataset contained 68,657 cells with median 1,732 genes and 2,963 transcripts per cell (median mitochondrial fraction 8.1%); the MCF10A dataset contained 113,720 cells with median 2,113 genes and 3,651 transcripts per cell (median mitochondrial fraction 10.7%).

**Gene quantification.** Count matrices were log-normalized (sc.pp.normalize_total to 10⁴ counts per cell followed by sc.pp.log1p). The full gene set is retained as the .raw attribute (NDF: 21,983 genes; MCF10A: 21,318 genes) and a highly variable gene subset was used for dimensionality reduction (NDF: 4,002 HVGs; MCF10A: 2,551 HVGs).

**Data files.** Primary processed data are stored as AnnData (h5ad) files at the locations: NDF 00_source_data/NDF/adata_NDF_after_diffexp.h5ad (2.1 GB) and MCF10A 00_source_data/MCF/adata_MCF_after_diffexp_v2.h5ad (4.4 GB). Network-entropy and Shannon-entropy derivatives used in Figure 7 are in 02_scent/outputs/{NDF,MCF10A}_scent.h5ad and 03_shannon/outputs/{NDF,MCF10A}_scent_shannon.h5ad. Raw sequencing reads will be deposited at NCBI GEO accession (REF) upon publication.

**Software.** All single-cell analyses used Scanpy v1.10 with default parameters except where noted. Module scoring used sc.tl.score_genes with random_state=42 and curated gene lists provided in Dataset S1. Differential expression used sc.tl.rank_genes_groups (Wilcoxon rank-sum test). UMAP embeddings used 50 principal components and 15 nearest neighbors; Leiden clustering used resolution 1.0.
